# Humoral immune responses against seasonal coronaviruses predict efficiency of SARS-CoV-2 spike targeting, FcγR activation, and corresponding COVID-19 disease severity

**DOI:** 10.1101/2021.09.14.460338

**Authors:** Jose L. Garrido, Matias Medina, Felipe Bravo, Sarah McGee, Francisco Fuentes, Mario Calvo, James W. Bowman, Christopher D. Bahl, Maria Inés Barría, Rebecca A. Brachman, Raymond A. Alvarez

## Abstract

Despite SARS-CoV-2 being a “novel” coronavirus, several studies suggest that detection of anti-spike IgG early in infection may be attributable to the amplification of humoral memory responses against seasonal hCoVs in severe COVID-19 patients. In this study, we examined this concept by characterizing anti-spike IgG from a cohort of non-hospitalized convalescent individuals with a spectrum of COVID-19 severity. We observed that anti-spike IgG levels positively correlated with disease severity, higher IgG cross-reactivity against betacoronaviruses (SARS-CoV-1 and OC43), and higher levels of proinflammatory Fc gamma receptor 2a and 3a (FcγR2a & FcγR3a) activation. In examining the levels of IgG targeting betacoronavirus conserved and immunodominant epitopes versus disease severity, we observed a positive correlation with the levels of IgG targeting the conserved S2’FP region, and an inverse correlation with two conserved epitopes around the heptad repeat (HR) 2 region. In comparing the levels of IgG targeting non-conserved epitopes, we observed that only one of three non-conserved immunodominant epitopes correlated with disease severity. Notably, the levels of IgG targeting the receptor binding domain (RBD) were inversely correlated with severity. Importantly, targeting of the RBD and HR2 regions have both been shown to mediate SARS-CoV-2 neutralization. These findings show that, aside from antibody (Ab) targeting of the RBD region, humoral memory responses against seasonal betacoronaviruses are potentially an important factor in dictating COVID-19 severity, with anti-HR2-dominant Ab profiles representing protective memory responses, while an anti-S2’FP dominant Ab profiles indicate deleterious recall responses. Though these profiles are masked in whole antigen profiling, these analyses suggest that distinct Ab memory responses are detectable with epitope targeting analysis. These findings have important implications for predicting severity of SARS-CoV-2 infections (primary and reinfections), and may predict vaccine efficacy in subpopulations with different dominant antibody epitope profiles.

## Introduction

The current coronavirus disease 2019 (COVID-19) pandemic is caused by the severe respiratory syndrome coronavirus 2 (SARS-CoV-2); which, to date, has infected more than 225 million and killed over 4.6 million people worldwide [1]. To combat the ongoing threat posed by SARS-CoV-2, researchers have made unprecedented progress in the last year and developed several antibody-based therapeutics and vaccines. While these therapies and vaccines are beginning to curtail the frequency and severity of SARS-CoV-2 infections, emerging variants continue to threaten the progress that has been made and reduce the efficacy of the current monoclonal antibodies (Ab) therapies and vaccines. Therefore, a better understanding of the Ab-mediated immune responses that correlate with protective responses could yield insights into the design of more effective, broadly-acting, antibody-based therapeutics and/or vaccines that target evolving viral variants.

Infection with SARS-CoV-2 has been observed to cause a wide range of disease severity. Generally, approximately 80% of individuals experience asymptomatic or mild disease, 15% experience moderately severe pneumonia, and 5% require hospitalization with severe acute respiratory distress syndrome (ARDS)[2]. Studies conducted in patients with severe COVID-19 commonly observed a hyper-immune activation state, which is characterized by highly elevated levels of cytokines, such as IL-6, TNF-α, and IL-8 [3-5]. This hyperimmune activation occurs 1-2 weeks after the onset of symptoms and coincides with the onset of adaptive immunity. Several studies have correlated early detection and higher IgG titers with more severe COVID-19 [6-12]. Specifically, studies have observed that elevated levels of antibodies targeting the spike protein correlate with severe hospitalized COVID-19 [12-16], suggesting that humoral immunity may be exacerbating the severity of COVID-19 [17-19]. Another correlate of severe COVID-19 and hyperimmune activation is the activation of monocytes and macrophages [20, 21], which have been hypothesized to contribute to hyperimmune activation through the release of high levels of proinflammatory cytokines [17, 22, 23]. Notably, Abs from severe COVID-19 patients have been shown to induce high levels of cytokine release from monocytes and macrophages *in vitro* [24, 25]. Altogether, these findings led several researchers to hypothesize that the onset of adaptive antibody immunity may be exacerbating COVID-19 severity through antibody dependent enhancement (ADE).

ADE effects have been observed in the context of several viral infections, including Dengue, RSV, and even other coronaviruses [26-28]. In general, ADE comes about due to Ab-mediated increases in virus replication and/or the level of proinflammatory mediators, thereby enhancing the severity of disease. In the case of dengue, primary infection with one of four dengue serotypes generally does not cause dengue hemorrhagic fever (DHF) and hyperimmune activation [27, 29]. However, upon secondary infection with a heterologous serotype, cross-reactive humoral memory responses can be activated and expanded, leading to inefficient IgG targeting of the secondary strain, resulting in DHF [30-32]. The hallmarks of DHF include elevated cytokine expression and viral titers, as well as high anti-dengue IgG titers; all of which have been associated with hyperimmune activation in severe COVID-19.

Studies examining the mechanisms underlying dengue-induced ADE have identified IgG-FcγR interactions as one of the primary factors governing increased disease severity [29, 33-35]. Fcγ Receptors (FcγRs) are a family of IgG-binding receptors expressed on the surface of innate immune effector cells (i.e., natural killer cells (NK), macrophages, monocytes, and Dendritic cells (DCs)) [36, 37]. In the context of viral infections, IgG-antigen immune complexes trigger a diverse array of non-neutralizing effector functions mediated through FcγR activation. These cellular effector functions include Ab-dependent cellular-cytotoxicity (ADCC), Ab-dependent cellular phagocytosis (ADCP), DC maturation and antigen presentation, and effector cytokine production [38-41]. Despite numerous studies demonstrating the critical importance of FcγR activation in controlling viral infections *in vivo* [42-46], in the context of viral infections that induce ADE effects, such as in DHF, high non-neutralizing IgG titers have been shown to be detrimental. The elevated levels of immune complexes exacerbate disease by facilitating the entry of virus particles into otherwise non-permissive host cells via FcγR, thereby increasing the kinetic and absolute levels of virus replication. Moreover, the interactions of these immune complexes with FcγR can also induce the production of proinflammatory cytokines, which at high levels increase localized tissue damage and mediate the most devastating effects attributed to “cytokine storms”, such as lung failure and cardiac arrest.

Pertinently, ADE effects have been reported with SARS-CoV-1 infections. *In vitro* viral replication assays performed using SARS-CoV-1 in the presence of anti-spike Abs show that Abs increase virus entry into B cells, monocytes, and macrophages via FcγR-IgG interactions [47-49], with some studies showing elevated induction of proinflammatory cytokines [13, 50]. Despite the enhanced uptake into cells, primary monocytes and macrophages fail to become productively infected, and replication has only been observed in primary B cells. Importantly, a vaccine study in hamsters demonstrated that despite the FcγR-IgG-mediated cellular uptake of SARS-CoV-1, vaccinated animals were protected in viral challenge experiments [51]. However, other *in vivo* studies in macaque models of SARS-CoV-1 infection have shown that anti-spike Abs increase the degree of acute lung injury in association with vaccine-induced or passively-infused Abs [13, 52]. Therefore, it still remains unclear whether anti-spike Abs definitively cause ADE responses *in vivo*, but some evidence suggests a potential role for anti-spike Abs in contributing to the elevated proinflammatory environment observed during severe SARS-CoV-1 infections in humans.

In most instances of virus-induced ADE, memory responses from a primary infection become activated and mount inefficient and detrimental responses to a similar, but distinct, heterologous strain, thereby interfering with evolution of *de novo* humoral responses against the contemporaneous strain. In general, *de novo* humoral responses are mounted by IgM, as it doesn’t require “class switching”, whereas IgG is part of a delayed secondary response. However, in the case of reinfection with a previously encountered pathogen, memory IgG responses can arise very quickly after antigen specific reactivation. Consistent with a recall response, SARS-CoV-2 specific IgG have been detected as early as 4-7 days after the onset of COVID-19 symptoms [7, 9, 11, 12], unlike the 7-14 days expected with a *de novo* antigen responses. Moreover, in some cases, IgG have been detected prior to, or contemporaneous with, IgM responses [6, 8].

While SARS-CoV-2 has now evolved variant strains that could potentially act like heterologous strains, these strains only arose several months after the initial outbreak and cannot account for any ADE-like effects observed early on in severe COVID-19 patients. In contrast, the seroprevalence of seasonal human coronaviruses (hCoVs) is fairly ubiquitous [53], with some seasonal hCoVs sharing regions of high sequence identity with SARS-CoV-2. In particular, the spike protein of SARS-CoV-2, which is the predominant viral antigen targeted by neutralizing Abs, contains regions of high sequence conservation, particularly in the S2 subunit. Notably, the S2 subunit contains the structural domains required for viral fusion [54]. In contrast, the RBD-containing S1 subunit bears far less sequence similarity with seasonal hCoVs. Importantly, the RBD region interacts with host ACE2 receptors, mediating viral attachment, fusion, and entry into cells, and is therefore the predominant site against which neutralizing Abs have been found to be directed in convalescent individuals [55-57]. Indeed, recent studies have found that SARS-CoV-2 naïve individuals possess IgG directed against the S2, but not the S1, subunit of the spike protein [58-60]. Additionally, some pre-pandemic serum samples were also found to have reactivity to the S2 subunit of the spike [58, 61]. Interestingly, pre-pandemic samples that displayed high SARS-CoV-2 reactivity also displayed high reactivity to the spike protein of OC43, a seasonal hCOV, and were found to be non-neutralizing against SARS-CoV-2 [58]. These findings show that IgG targeting of the S2 region exists prior to SARS-CoV-2 infection in some individuals, with the potential for these responses to become amplified upon SARS-CoV-2 infection. Indeed, some studies have already observed that cross-reactive Ab responses against OC43 spike protein become elevated in SARS-CoV-2 convalescent individuals [10, 58, 62]. However, the contribution of seasonal hCoV cross-reactive IgG to the humoral response against SARS-CoV-2 remains unclear, with some studies showing no correlation with severity [63], while others observed a correlation with severity [10, 62].

Beyond the broad characterization of IgG S1 vs S2 binding, other studies have conducted peptide walks to more precisely identify the linear epitope hotspots targeted by humoral responses against the spike protein. These studies have identified several immunodominant regions which overlap or flank several functional features in the SARS-CoV-2 spike protein, such as the S1/S2 junction, S2’ fusion peptide site, and the HR1 and HR2 sites [64-67]. When these immunodominant regions were examined in relation to COVID-19 severity, several epitopes were highly correlated with severity [64, 66]. Interestingly, some epitopes were also highly conserved amongst seasonal coronaviruses [62, 68]. These findings, coupled with IgG-recognition of the S2 region in naïve individuals and the elevated levels of OC43 cross-reactive IgG in convalescent individuals, suggest that preexisting recall responses against seasonal coronaviruses likely contribute to the humoral response against SARS-CoV-2. However, the contribution of these cross-reactive Abs to the evolution of effective humoral immunity, disease severity, and outcomes remains unclear; previous research has failed to establish a conclusive, significant contribution—either protective or deleterious. However, these studies either relied on pooled analyses, which masks complex signatures, or did not examine disease severity. Therefore, there is an immediate need to characterize the FcγR activation and Ab targeting profiles associated with mild versus severe SARS-CoV-2 infections. Additionally, there is also a need to understand whether cross-reactive recall responses targeting non-neutralizing regions could be inducing inefficient and/or ADE-like effects that are detrimental for COVID-19 outcomes. In this study, we investigate how anti-spike IgG responses correlate with disease severity, FcγR activation, and epitope-targeting in a cohort of non-hospitalized SARS-CoV-2 convalescent donors, as well as dissect the contributions of divergent recall responses against seasonal coronaviruses to the SARS-CoV-2 humoral response.

## Results

### Non-hospitalized SARS-CoV-2 convalescent individuals displayed a spectrum of COVID-19 symptom severity

In order to obtain a better understanding of the humoral responses generated by SARS-CoV-2 convalescent individuals, a total of 48 donors were recruited in the New York city area during the first wave of the COVID-19 pandemic in the spring of 2020. The cohort was separated into two groups based on the history of a positive or negative SARS-CoV-2 test (PCR or serology): convalescent donors (positive test; CONVALESCENT) and negative controls (negative test; NAÏVE). The negative control group was composed of age- and sex-matched SARS-CoV-2 naïve donors (Table 1). Importantly, in convalescent donors, the median time from the onset of symptoms to the time of blood draw was 43 days, ensuring that anti-SARS-CoV-2 IgG levels were sufficiently high for pathogen-specific IgG analysis in all convalescent donors.

**Table 1.**
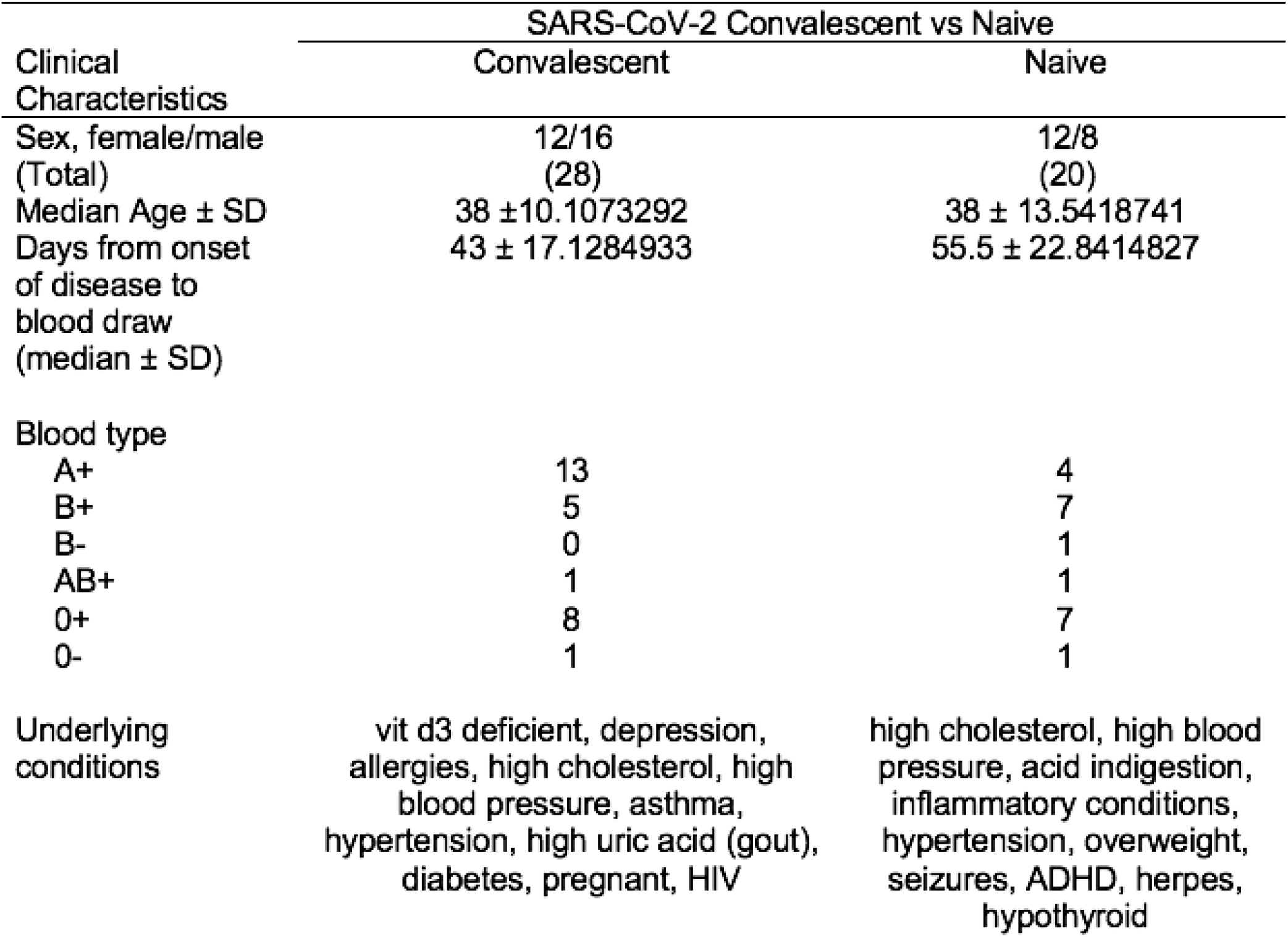
Characteristics of SARS-CoV-2 convalescent and naïve donors.

| Clinical Characteristics | SARS-CoV-2 Convalescent vs Naive |  |
| --- | --- | --- |
|  | Convalescent | Naive |
| Sex, female/male (Total) | 12/16 (28) | 12/8 (20) |
| Median Age ± SD | 38 ± 10.1073292 | 38 ± 13.5418741 |
| Days from onset of disease to blood draw (median ± SD) | 43 ± 17.1284933 | 55.5 ± 22.8414827 |
| Blood type |  |  |
| A+ | 13 | 4 |
| B+ | 5 | 7 |
| B- | 0 | 1 |
| AB+ | 1 | 1 |
| O+ | 8 | 7 |
| O- | 1 | 1 |
| Underlying conditions | vit d3 deficient, depression, allergies, high cholesterol, high blood pressure, asthma, hypertension, high uric acid (gout), diabetes, pregnant, HIV | high cholesterol, high blood pressure, acid indigestion, inflammatory conditions, hypertension, overweight, seizures, ADHD, herpes, hypothyroid |

To assess the relative severity of disease, we surveyed the symptomological history of disease for each convalescent donor, including the intensity and duration of each symptom. This information was used to calculate both an average severity score for each symptom (sup Table 2), along with a composite symptom severity score for every convalescent donor. In doing so, we observed that the convalescent group presented a wide range of symptoms and severities ranging from mild to more severe COVID-19. The most commonly reported symptoms were cough, headache, fatigue and myalgia, with greater than 80% of the cohort experiencing these symptoms. The symptoms with the highest average scores were fatigue, myalgia, diarrhea, and fever. The least frequently experienced symptoms were nausea and vomiting. In calculating a composite symptom severity score for each convalescent individual, we observed that convalescent donors clustered into two categories: those that experienced milder symptoms (n=13), with composite severity scores below 45; and those with more severe disease (n=15), with scores above 45. For simplicity, these two groups are henceforth referred to as MILD and SEVERE, respectively, as they reflect these two ends of the non-hospitalized COVID-19 spectrum—with the acknowledged caveat that all donors represent non-hospitalized, non-fatal COVID-19 cases. Despite not being hospitalized, individuals with the highest severity scores (>45) commonly experienced high fevers (>101°C) for more than a week, severe myalgia, headaches, and difficulty breathing. Additionally, two severe patients experienced weight loss of >15% of body weight. Difficulty breathing was the most differentially experienced symptom in severe versus mild infections, with 60% of severe donors experiencing this symptom versus only 15% of mild donors (Table 2). The only other symptom that was differentially reported amongst mild versus severe donors was diarrhea, with approximately 33% of severe donors experiencing this symptom versus 15.4% of mild donors. For all other symptoms, there was a comparable frequency of donors that experienced each symptom, with the main difference being intensity.

**Table 2.** Symptomology of SARS CoV-2 convalescent donors.

|  | All convalescent donors |  |  | Mild convalescent donor |  |  | Severe convalescent donor |  |  |
| --- | --- | --- | --- | --- | --- | --- | --- | --- | --- |
| Symptoms | Frequency of symptoms (N=28) | Average symptom severity score (1-10) | Average Duration of Symptoms (Days) | Frequency of symptoms (N=13) | Average symptom severity score (1-10) | Average Duration of Symptoms (Days) | Frequency of symptoms (N=15) | Average symptom severity score (1-10) | Average Duration of Symptoms (Days) |
| Runny Nose | 14/28 (50.0%) | 5.14 | 9.20 | 7/13 (53.9%) | 3.43 | 8.00 | 7/15 (46.7%) | 6.86 | 8.67 |
| Sore throat | 16/28 (57.1%) | 5.06 | 5.25 | 6/13 (53.8%) | 3.75 | 3.50 | 10/15 (66.7%) | 5.70 | 4.60 |
| Cough | 21/28 (75.0%) | 5.33 | 8.50 | 9/13 (69.2%) | 4.00 | 6.43 | 12/15 (80%) | 6.33 | 9.00 |
| Fatigue | 25/28 (89.3%) | 6.56 | 7.24 | 12/13 (92.3%) | 4.36 | 4.73 | 13/15 (86.7%) | 8.39 | 10.0 |
| Myalgia | 24/28 (85.7%) | 6.83 | 6.33 | 11/13 (84.6%) | 4.46 | 3.40 | 13/15 (86.7%) | 8.62 | 8.69 |
| Diarrhea | 7/28 (25.0%) | 6.14 | 8.14 | 2/13 (15.4%) | 5.50 | 2.00 | 5/15 (33.3%) | 6.40 | 9.80 |
| Fever | 19/28 (67.9%) | 6.11 | 4.47 | 9/13 (69.2%) | 4.09 | 2.78 | 10/15 (66.7%) | 7.70 | 6.22 |
| Vomitting | 5/28 (17.9%) | 4.30 | 4.00 | 3/13 (23.1%) | 4.00 | 4.00 | 2/15 (13.3%) | 4.75 | 3.00 |
| Nausea | 4/28 (14.3%) | 5.50 | 4.00 | 2/13 (15.4%) | 4.50 | 6.00 | 2/15 (13.3%) | 6.50 | 7.00 |
| Headache | 24/28 (85.7%) | 5.50 | 7.69 | 11/13 (86.6%) | 3.85 | 3.70 | 13/15 (86.7%) | 6.77 | 10.0 |
| Shortness of breadth | 11/28 (39.3%) | 4.00 | 8.00 | 2/13 (15.4%) | 3.00 | 2.00 | 9/15 (60%) | 4.22 | 8.11 |

Table depicts frequency of COVID-19 symptoms experienced by convalescent donors. Symptom intensity was scored out of 10, with 10 being most severe, and 0 not being experienced. Mild convalescent donors were defined as donors with composite symptom intensity scores below 45. Severe convalescent donors were defined as donors with composite symptom intensity scores above 45.

### COVID-19 severity correlates with higher anti-spike Ab titers in SARS-CoV-2 convalescent individuals

To elucidate the humoral immune responses associated with mild to severe COVID-19, we began by quantifying the levels of IgG directed against the SARS-CoV-2 spike protein in convalescent donors. To quantify the levels of anti-spike IgG in convalescent donors, we first utilized a conventional ELISA-based assay [69]. Using this assay, we detected significantly higher levels of anti-spike IgG titers in the convalescent versus SARS-CoV-2 naïve donors, with only a few naïve donors possessing titers above background (Figure 1A). Amongst the convalescent donors, we observed a range of anti-spike IgG levels, which differed approximately 30-fold between lowest and highest samples (Figure 1B). Interestingly, using conventional ELISA, two donors that tested PCR-positive for SARS-CoV-2 infection possessed no detectable anti-spike titers and eight additional convalescent donors possessed titers that were less than 3-fold above background, which was comparable to the anti-spike levels observed in some SARS-CoV-2 naïve donors. We next compared the anti-spike IgG levels against the composite severity scores of each donor and observed a significant positive correlation between anti-spike titers and higher severity scores (Figure 1C). Importantly, several studies have found similar correlations between anti-spike IgG and COVID-19 severity in hospitalized patients [6-9, 11, 12].

**Figure 1.**
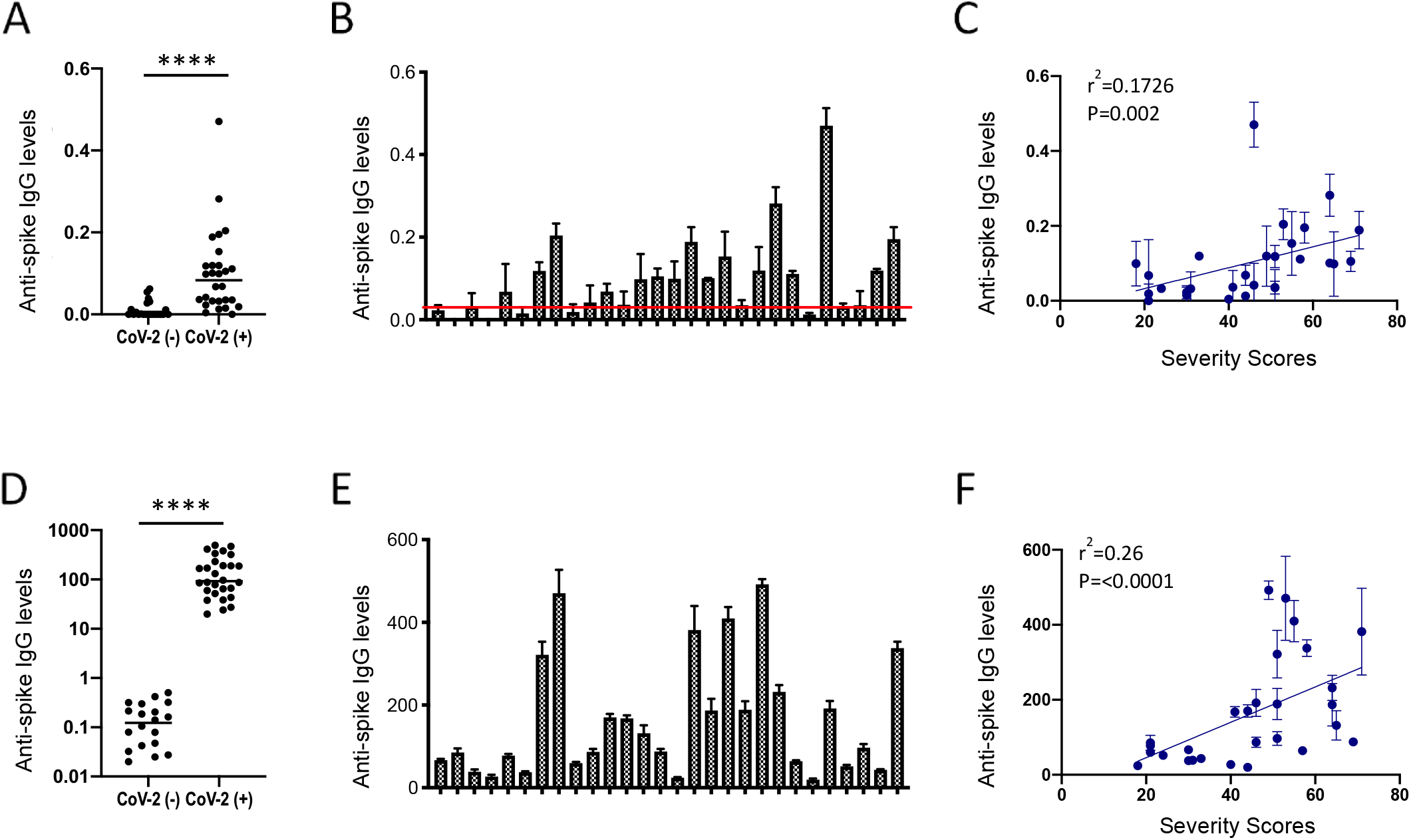
Anti-spike IgG levels in convalescent donors correlates with COVID-19 severity. **(A)** Anti-spike IgG titers in SARS-CoV-2 naïve (CoV-2-) and SARS-CoV-2 positive convalescent (CoV-2+) donors, as quantified by recombinant spike protein ELISA. **(B)** Variations in anti-spike titers among convalescent (CoV-2+) donors as quantified by ELISA. Red line represents 3-fold above the mean anti-spike levels of naïve (CoV-2-) donors, as quantified by ELISA. **(C)** Comparison of anti-spike IgG titers, as quantified by ELISA, vs. composite severity scores amongst convalescent donors. **(D-F)** Levels of anti-spike IgG in naïve (CoV-2-) and convalescent (CoV-2+) donors as quantified using a cell-based assay. **(D)** Anti-spike IgG titers in naïve (CoV-2-) and convalescent (CoV-2+) donors as quantified by cell-based assay. **(E)** Variations in anti-spike titers amongst convalescent (CoV-2+) donors as quantified by cell-based IgG binding assay. **(F)** Comparison of anti-spike IgG titers, as quantified by ELISA, vs. composite severity scores amongst convalescent donors. The SEM of N=3 experiments are shown.

In addition to neutralization, IgG also mediates non-neutralizing effector responses against virus-infected host cells, which involve the recognition of viral antigens expressed on the surface of host cells [38-41]. Therefore, we next quantified the levels of anti-spike IgG binding to cell surface-expressed forms of the SARS-CoV-2 spike. To quantify cell surface IgG binding, we employed a cell-based system using 293T endothelial cells transfected to express the SARS-CoV-2 spike. In this system, the levels of anti-spike IgG are quantified using a binding index that accounts for both the percentage and Median Fluorescence Intensity (MFI) of IgG bound to spike-expressing cells. We have previously used this method to discern viral antigen-specific IgG levels in convalescent donors [70-72]. The benefit of this composite metric is that both frequency and density of IgG-antigen binding are captured, as both parameters contribute to the IgG-mediated activation of non-neutralizing effector functions. Additionally, more than in ELISA, expressing spike protein in a human cell-based assay likely retains tertiary structure and glycosylation consistent with natural infection, potentially capturing a greater range of paratopes (i.e., higher sensitivity).

Using this cell-based assay, we detected an approximate 1.5-log difference in anti-spike IgG titers between naïve and SARS-CoV-2 convalescent donors, and a further 2-log difference amongst the convalescent donors (Figure 1D, 1E). Interestingly, the two PCR-positive SARS-CoV-2 donors whose anti-spike IgG levels were undetectable by ELISA had detectable, albeit low, anti-spike IgG levels in the cell-based assay. In the cell-based assay, these levels were 1 log above the values obtained with IgG from naïve donors, demonstrating greater resolution as compared to ELISA. Additionally, using the cell-based assay, anti-spike IgG levels were readily detectable above negative control (naïve) IgG levels in the eight donors that possessed low anti-spike IgG levels not clearly discernible above background using the ELISA assay, demonstrating the high sensitivity and specificity of this cell-based assay (Supplementary Figure 1A). Similar to the ELISA results, we observed a strong positive correlation between anti-spike IgG levels and disease severity (r^2^ = 0.27 p < 0.0001) (Figure 1F), with severe donors possessing significantly higher anti-spike IgG levels (Supplementary Figure 1B).

### Higher levels of anti-spike IgG-mediated FcγR activation corelates with COVID-19 severity

Elevated levels of anti-spike IgG and proinflammatory cytokine levels are detected in severe hospitalized COVID-19 patients, suggesting that IgG maybe exacerbating disease severity via FcγR mediated ADE effects. Therefore, we next examined the levels of FcγR-signaling induced by IgG from convalescent individuals. Specifically, we examined the levels of Fc-γ receptors (FcγR), FcγR2a and FcγR3a, since these FcγR activate several important cellular effector functions, like ADCC and ADCP [36, 37]. Importantly, both of these FcγR are capable of activating an array of pro-inflammatory cytokines.

To quantify the ability of patient sera to activate FcγR, we transfected 293T cells to express the SARS-CoV-2 spike. The spike-expressing 293T cells were co-cultured with FcγR2a or FcγR3a reporter cell lines in the presence of various concentrations of purified donor IgG [70, 71]. Using this system, we observed a wide range of FcγR activation in response to convalescent donor IgG, but only background levels of activation in response to SARS-CoV-2-naïve donor IgG (Figure 2A). In examining the relationship between disease severity and FcγR activation, we observed a strong positive correlation between severity scores and FcγR2a and FcγR3a activation (Figure 2B, 2D). This relationship was even more apparent in severe donors, as compared to mild donors (Figure 2A and 2C). Interestingly, the majority of IgG from mild donors did not induce greater FcγR2a-activation than SARS-CoV-2 naïve control IgG (Figure 2A). In contrast, the majority of IgG from severe donors induced FcγR2a signaling that was 2-fold above naïve control IgG. In regards to FcγR3a activation, both mild and severe convalescent donors induced signaling that was at least 2-fold above naïve control IgG (Figure 2C). Similar to FcγR2a-activation, IgG from severe donors possessed significantly higher FcγR3a activity as compared to IgG from mild donors.

**Figure 2.**
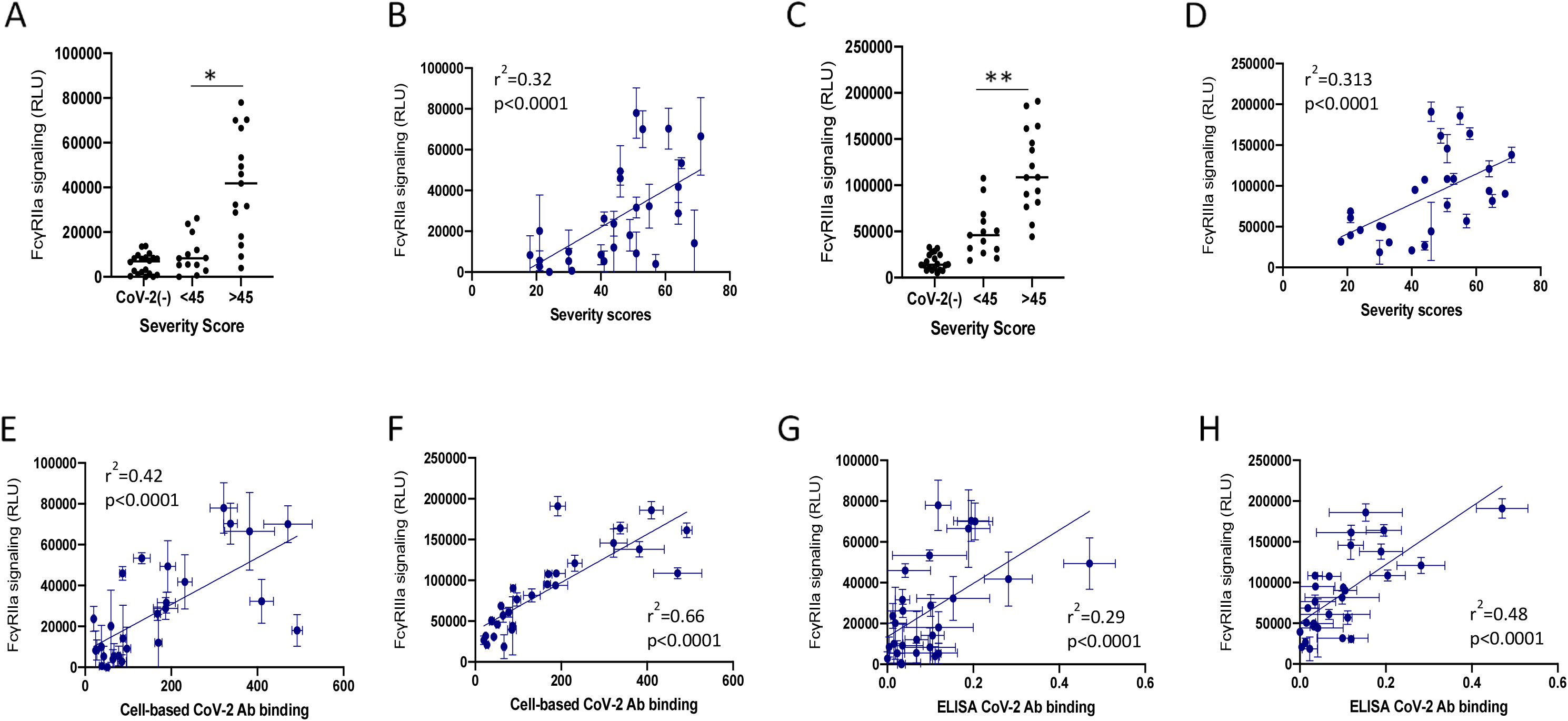
Levels of anti-spike lgG-induced FcγR-activation correlates with COVID-19 severity and anti-spike titers. Figure depicts the levels of FcγR-signaling induce by purified lgG derived from SARS-CoV-2 convalescent donors in response to SARS-CoV-2 spike protein expressed on the surface of 293T cells. Graphs show the levels of **(A**,**B)** FcγR2a and **(C**,**D)** FcγR3a signaling using 25 μg/ml lgG versus composite symptom severity scores. Graphs show comparison of donor groups separated by CoV-/+ status and COVID-19 severity scores versus the levels of **(B)** FcγR2a or **(D)** FcγR3a signaling using 25 μg/ml of lgG. Figures **E-H** compare the anti-spike lgG titers as quantified by cell-based or ELISA assays versus the levels of FcγR signaling induced by purified lgG derived from SARS-CoV-2 convalescent donors. Graphs show the level of anti-spike lgG as quantified by cell-based assay versus the levels of **(E)** FcγR2a and **{G)** FcγR3a signaling using 25 μg/ml lgG. Graphs show the level of anti-spike lgG as quantified by ELISA versus the levels of **(F)** FcγR2a and **(H)** FcγR3a signaling using 25 μg/ml lgG. All FcγR results are the SEM of N=3 experiments.

Next, we evaluated the relationship between FcγR-activation and the levels of anti-spike IgG. We observed a significant positive correlation with both FcγR2a and FcγR3a signaling and the levels of anti-spike IgG in all convalescent donors, as quantified by both ELISA and cell-based anti-spike IgG binding assays (Figure 2E-H). However, anti-spike IgG levels as quantified by the cell-based assay were much more highly correlated with the levels of FcγR3a signaling (R^2^= 0.66)(Figure 2F), as compared to anti-spike titers obtained by ELISA (R^2^= 0.48)(Figure 2H). In addition, FcγR3a signaling in general was much more directly correlated with anti-spike IgG levels (R^2^= 0.66; R^2^= 0.48)(Figure 2F, 2H), as compared to FcγR2a signaling (R^2^= 0.43; R^2^= 0.29)(Figure 2E, 2G). Altogether, these data show a strong positive correlation between anti-spike IgG titers, the levels of FcγR activation, and COVID-19 severity.

### COVID-19 disease severity is correlated with higher anti-spike IgG cross-reactivity against betacoronaviruses

Normally, IgG responses against novel antigens arise days to weeks after the establishment of infection, since naïve B cells require 7-14 days to become activated, class-switch, and begin producing IgG [73]. However, in the case of COVID-19, elevated IgG has been reported within 4 days of infection [7, 9, 11]. The seroprevalence of seasonal human alpha- and betacoronaviruses are ubiquitous [53], with 4-27% of the population testing positive for any given hCoV at any given time [61]. The spike proteins of these seasonal hCoVs share approximately 25% and 30% sequence identity with the spike protein of SARS-CoV-2, respectively (Table 3). The high seroprevalence of IgG against common coronaviruses, coupled with the high degree of identity between common seasonal hCoVs and SARS-CoV-2 spike, lead us to hypothesize that humoral responses against common coronaviruses may contribute to SARS-CoV-2 anti-spike IgG responses.

**Table 3.**
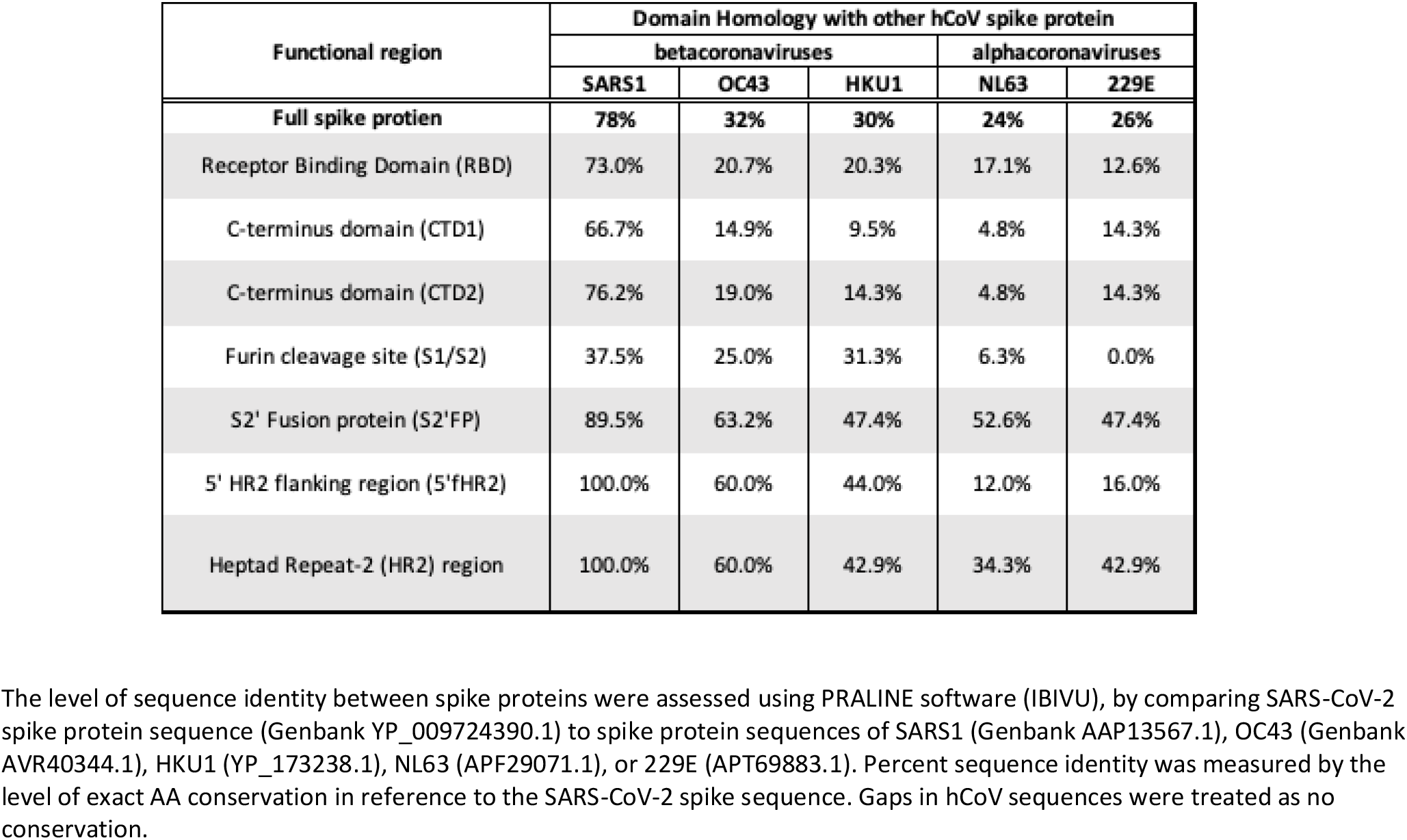
Sequence identity of SARS CoV-2 Immunodominant epitopes and functional regions in comparison to seasonal hCoVs.

| Functional region | Domain Homology with other hCoV spike protein |  |  |  |  |
| --- | --- | --- | --- | --- | --- |
|  | betacoronaviruses |  |  | alphacoronaviruses |  |
|  | SARS1 | OC43 | HKU1 | NL63 | 229E |
| Full spike protien | 78% | 32% | 30% | 24% | 26% |
| Receptor Binding Domain (RBD) | 73.0% | 20.7% | 20.3% | 17.1% | 12.6% |
| C-terminus domain (CTD1) | 66.7% | 14.9% | 9.5% | 4.8% | 14.3% |
| C-terminus domain (CTD2) | 76.2% | 19.0% | 14.3% | 4.8% | 14.3% |
| Furin cleavage site (S1/S2) | 37.5% | 25.0% | 31.3% | 6.3% | 0.0% |
| S2' Fusion protein (S2'FP) | 89.5% | 63.2% | 47.4% | 52.6% | 47.4% |
| 5' HR2 flanking region (5'fHR2) | 100.0% | 60.0% | 44.0% | 12.0% | 16.0% |
| Heptad Repeat-2 (HR2) region | 100.0% | 60.0% | 42.9% | 34.3% | 42.9% |

The level of sequence identity between spike proteins were assessed using PRALINE software (IBIVU), by comparing SARS-CoV-2 spike protein sequence (Genbank YP\_009724390.1) to spike protein sequences of SARS1 (Genbank AAP13567.1), OC43 (Genbank AVR40344.1), HKU1 (YP\_173238.1), NL63 (APF29071.1), or 229E (APT69883.1). Percent sequence identity was measured by the level of exact AA conservation in reference to the SARS-CoV-2 spike sequence. Gaps in hCoV sequences were treated as no conservation.

To examine this hypothesis, we measured the levels of anti-spike IgG binding against common hCoV spike proteins using IgG from SARS-CoV-2 convalescent and naïve donors. Specifically, we measured the amount of IgG binding to the spike of a seasonal betacoronavirus, OC43, as well as two alphacoronaviruses, NL63 and 229E. Additionally, we examined the level of cross-reactivity of SARS-CoV-2 convalescent IgG against the spike protein of SARS-CoV-1 (SARS1). Since we observed more sensitive detection of anti-spike IgG binding using the cell-based assay versus ELISA, we quantified the levels of seasonal hCoV anti-spike IgG binding using cell-based assays. Similar to SARS-CoV-2, cell-based IgG binding assays showed that naïve donor IgG had no recognition of SARS1 spike protein (Figure 3A). The majority of SARS-CoV-2 convalescent donors, however, did possess IgG which recognized the SARS1 spike protein. IgG-recognition of the OC43 spike was significantly elevated in SARS-CoV-2 convalescent donors as compared to SARS-CoV-2 naïve donors (Figure 3B). In contrast, IgG binding to the alphacoronavirus spike proteins NL63 and 229E was not significantly different amongst SARS-CoV-2 convalescent and naïve donors (Figure 3C, 3D). This finding suggests that IgG responses against SARS-CoV-2 did not boost IgG responses against alphacoronaviruses. When further explored in relation to disease severity, IgG from severe donors possessed higher cross-reactivity to the spike protein of SARS1 and OC43 as compared to mild donors (Figure 3E, 3F). These findings show that infection with SARS-CoV-2 induces IgG that are cross-reactive with other betacoronaviruses, and higher levels of cross-reactive IgG correlate with more severe disease.

**Figure 3.**
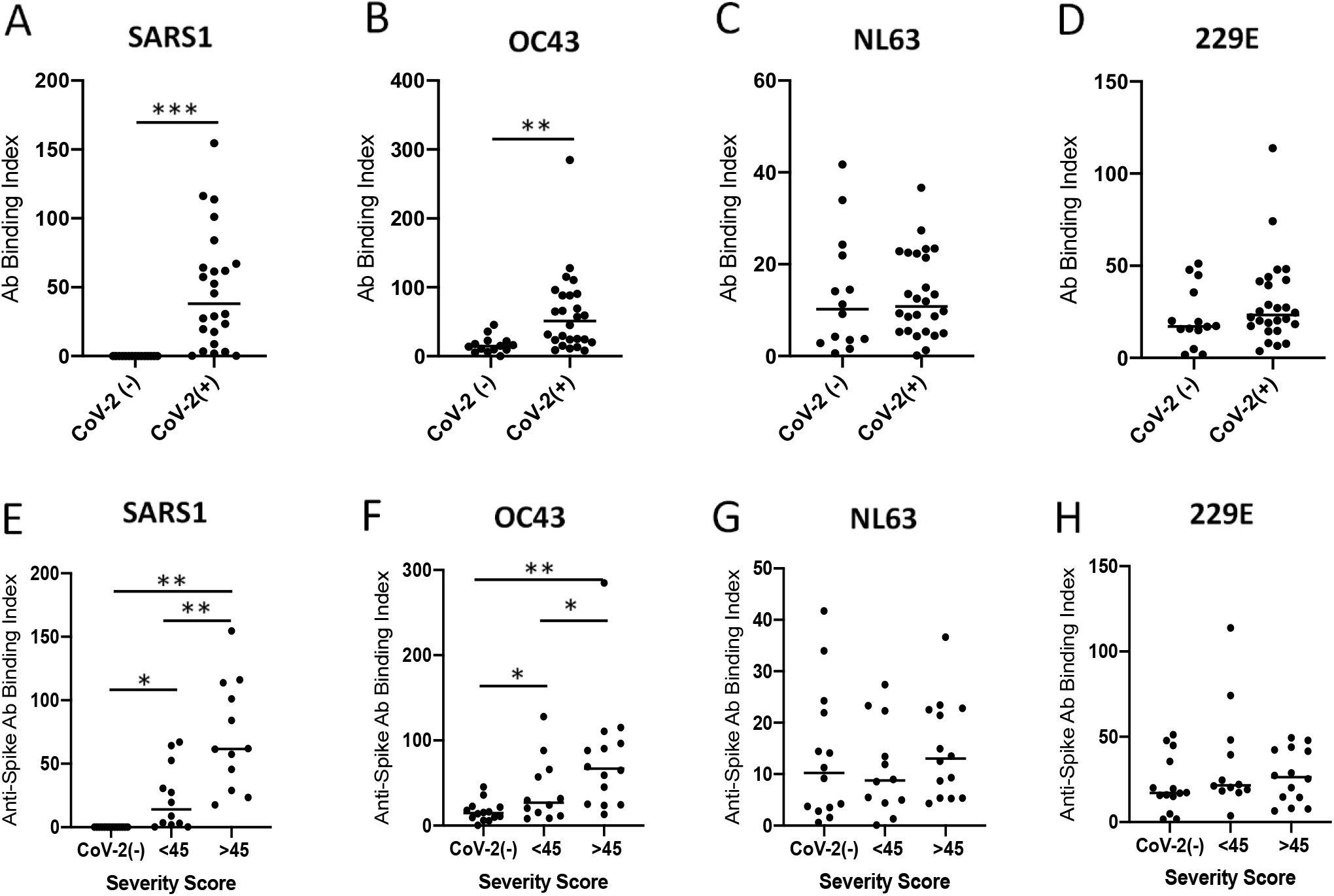
Higher titers of IgG against non-CoV-2 betacoronaviruses are correlated with COVID-19 severity. **(A-H)** Graphs compare the level of IgG binding to spike proteins of **(A**,**E)** SARS1, **(B**,**F)** OC43, **(C**,**G)** NL63 and **(D**,**H)** 229E coronavirus as assessed by cell-based assay and detected by flow cytometry. Graphs **A-D** compare the level anti-spike IgG in SARS-CoV-2 naïve versus convalescent donors. Graphs **E-H** compare the levels of anti-spike IgG among donor groups separated by SARS-CoV-2 status and COVID-19 severity scores. For all cross-reactive Ab binding results, the SEM of N=3 experiments are shown.

### Immunodominant regions with high OC43 sequence identity differentially correlate with COVID-19 severity

Recently, several groups have conducted SARS-CoV-2 peptide walk experiments using Abs from hospitalized COVID-19 patients to identify immunodominant epitopes that correlate with severe COVID-19 [64-67]. Notably, some of the immunodominant epitopes that have been identified overlap with functional regions within the SARS-CoV-2 spike protein, which include the S1/S2 furin cleavage site (S1/S2) and S2’ cleavage fusion protein region (S2’FP), as well as regions in and around heptad repeat 1 and 2 (HR1 & HR2). Interestingly, several of these regions share high sequence identity with seasonal coronaviruses [62, 68], suggesting that IgG-recognition of these immunodominant epitopes could be due to recall responses to seasonal coronaviruses such as OC43.

To explore how epitope-targeting relates to infection severity in non-hospitalized individuals, we screened our cohort against a panel of immunodominant SARS-CoV-2 peptides that possessed high identity with seasonal hCoVs (Table 3), in particular betacoronavirus OC43 (Figure 4). To contrast these conserved regions, we also screened the levels of IgG targeting immunodominant regions that possessed little identity with seasonal CoVs. Broadly, this set of peptides represent most of the functional regions within the spike protein, including S1/S2 furin cleavage site, S2’ cleavage site, and the HR2 region (Table 3). In addition, we quantified the levels of IgG targeting the RBD region, since several studies have identified that Abs targeting the RBD region mediate potent neutralization *in vitro* and control of SARS-CoV-2 infection *in vivo* [55-57, 74].

**Figure 4.**
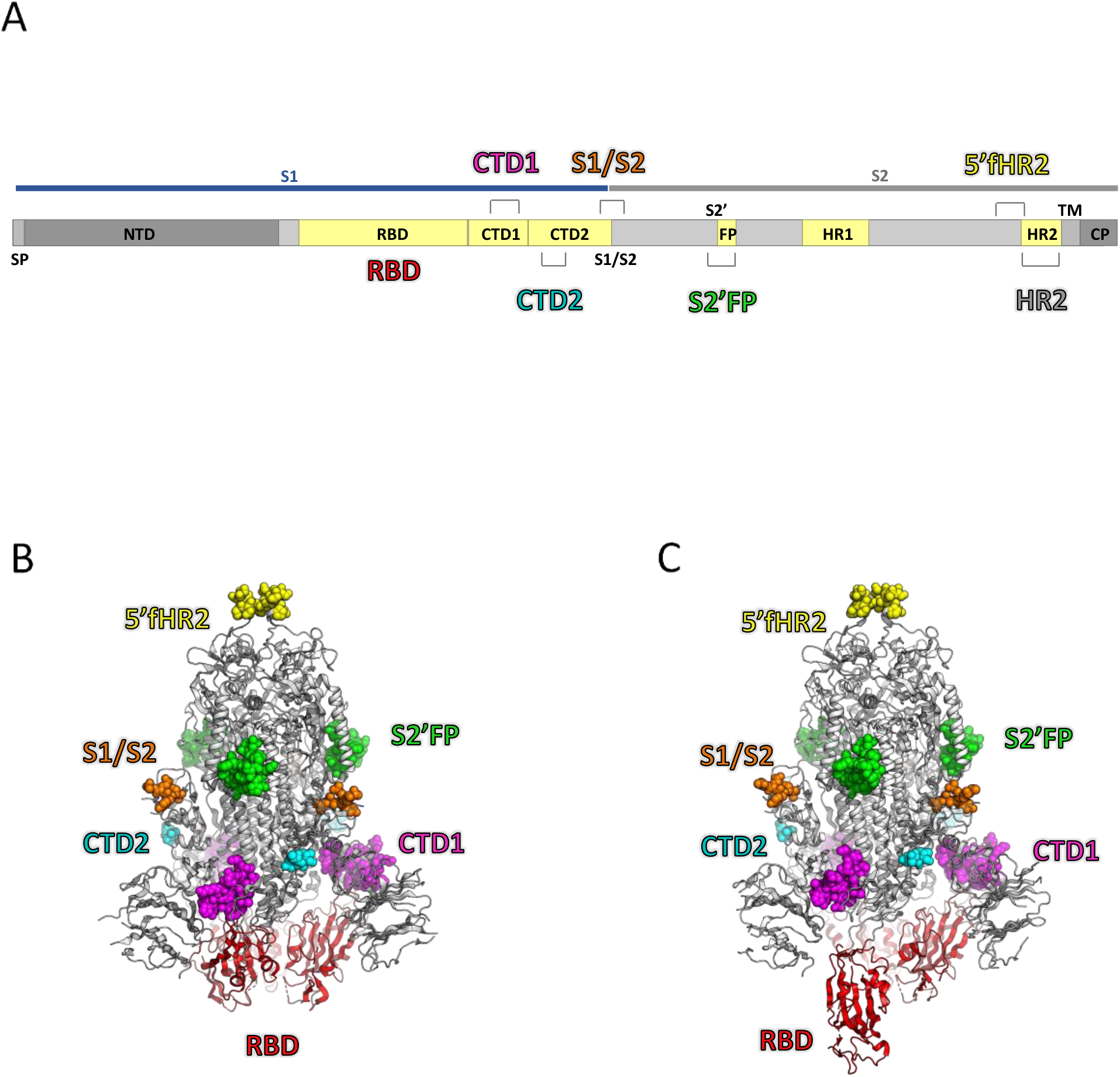
Localization of SARS CoV-2 spike immunodominant regions. **A)** Diagram depicts SARS CoV-2 spike protein subdomains, which include the N-terminal Domain (NTD); Receptor Binding Domain (RBD); S1-C-terminus Domains 1&2 (CTD1&2); Furin cleavage site (S1/S2); S2’ cleavage site & fusion protein domain (S2’FP); Heptad Repeat Domains 1&2 (HR1&2); transmembrane domain (TM) and cytosolic domain (CP). Immuno-dominant regions with either high or low sequence identity with betacoronavirus OC43 are shown, where an asterisk indicates 100% sequence identity (Red) and a consistency score of zero (Blue) indicates no conservation of amino acid characteristics. **(B**,**C)** The homotrimeric SARS-CoV-2 spike protein is shown in the (**B**) closed and (**C**) open state (PDB IDs 6VXX and 6VYB, respectively). In each protomer of the spike, the protein mainchain is shown as a cartoon representation and colored white, except for the RBD, which is colored red. The atoms in the six peptide epitopes that were tested are shown as space-filling models, colored according to peptide number. There are regions of missing density in the models, presumably due to conformational flexibility, and these regions are omitted here; CTD1 (magenta) and S2’FP (green) are fully resolved, CTD2 (cyan), S1/S2 (orange), and 5’fHR2 (yellow) are partially resolved, and HR2 is completely absent in the structure.

To quantify the levels of IgG targeting these regions, we developed a luciferase-based ELISA to allow for the sensitive detection of IgG binding. Using this assay, we first examined the levels of IgG targeting the RBD region. We observed a significant inverse correlation between the levels of anti-RBD IgG and overall severity scores among convalescent donors (P=<0.0001)(Figure 5A). In comparing donors with milder versus severe disease, we observed significantly higher levels of anti-RBD IgG-targeting in donors with mild disease (sup Figure 3A). Notably, no significant correlation was observed between anti-RBD IgG and total anti-spike IgG levels (Figure 5B).

**Figure 5.**
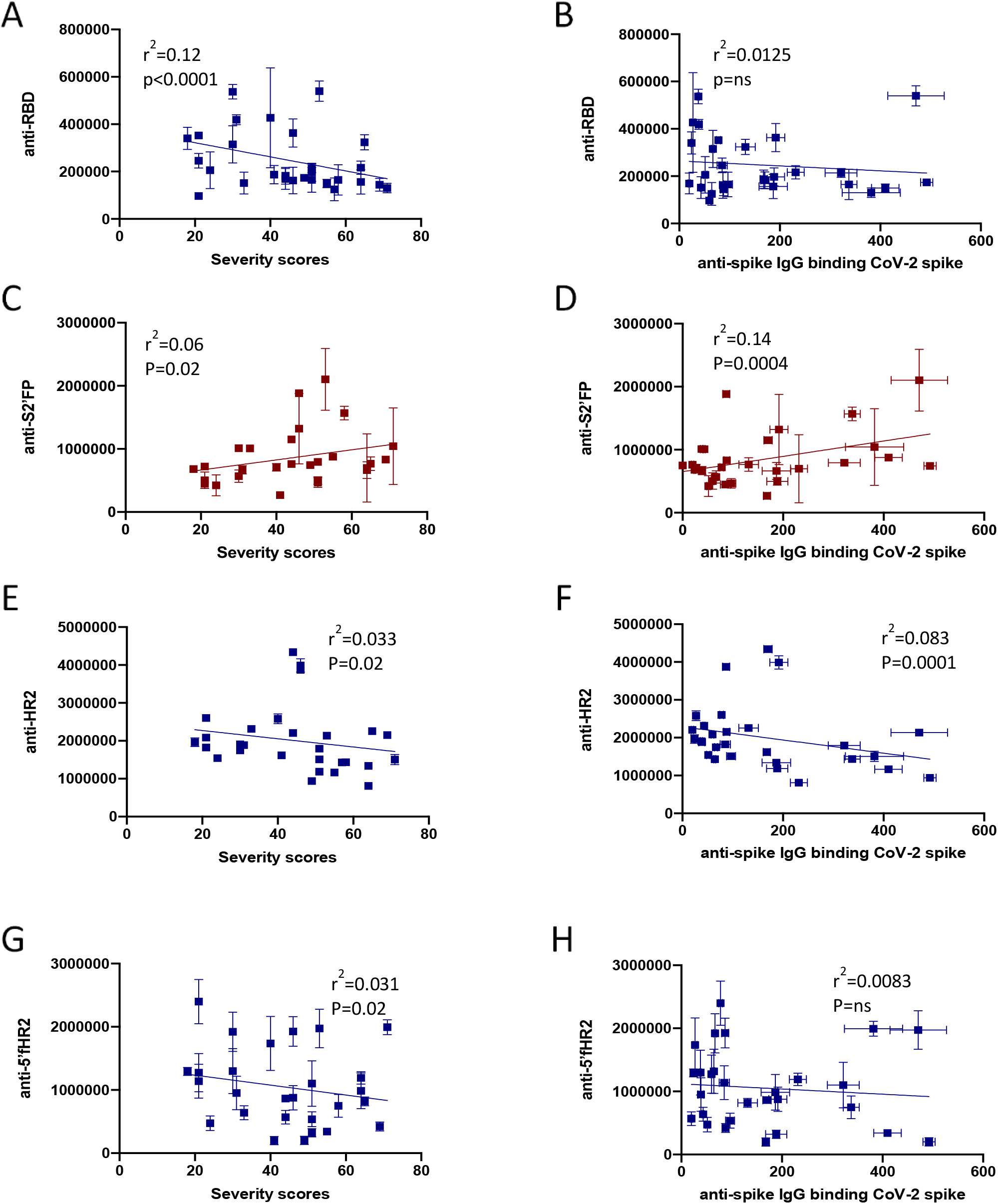
SARS-CoV-2 convalescent IgG differentially target seasonal CoV-conserved SARS-CoV-2 immunodominant epitopes. Graphs compare the levels of IgG-binding to **(A**,**B)** RBD, **(C**,**D)** S2’FP, **(E**,**F)** HR2, **(G**,**H)** 5’F HR2 regions as detected by luminescent ELISA versus **(A, C, E, G)** severity scores or **(B, D, F, H)** total anti-spike levels quantified by cell-based binding assay. The levels of IgG targeting are the SEM of N=3 experiments are graphed.

We next quantified the levels of IgG-targeting of three representative immunodominant peptides with high sequence identity to seasonal hCoVs. One selected peptide contains the immunodominant region of S2’FP and possesses the highest sequence identity with OC43 among the screened immunodominant regions. The fusion protein (FP) region becomes exposed after cleavage of the spike protein, which induces conformational rearrangement in the spike allowing for insertion of the FP into the host cell membrane, thereby facilitating viral fusion [75, 76]. We observed a significant positive correlation between the levels of IgG binding the S2’FP region and severity (Figure 5C). We also observed that IgG-targeting of this region increased with anti-spike IgG titers (Figure 5D). Next, we examined the levels of IgG binding to the Heptad Repeat 2 (HR2) domain. HR2 functions to mediate viral fusion and entry into host cells through the formation of a six-helix bundle in conjunction with the HR1 domain [77]. In contrast to IgG targeting the S2’FP region, we observed a significant inverse correlation with the levels of IgG targeting the HR2 region and severity (Figure 5E); on average, mild donors possessed higher levels of IgG targeting the HR2 region, as compared to severe donors (sup Figure 3E). We also observed that IgG-targeting of the HR2 region was inversely correlated with overall anti-spike IgG titers (Figure 5F). Lastly, we examined IgG-targeting of an immunodominant and hCoV conserved region located just upstream of the HR2 domain, aka the 5’ flank HR2 (5’fHR2). The levels of IgG binding in this region was also inversely correlated with severity (Figure 5G), similar to direct targeting of the HR2 region; however, IgG levels were not inversely correlated with overall anti-spike IgG titers (Figure 5H).

To contrast and compare the conserved region targeting data, we next examined the levels of IgG targeting immunodominant regions that were not highly conserved with OC43 or any other seasonal hCoV (Figure 6). One region we examined contained the novel furin cleavage site at the S1/S2 junction, while the other two regions were located within the C-terminal domain (CTD) just downstream of the RBD region (Figure 4). In screening these regions, we observed a significant positive correlation with the levels of CTD1 targeting and severity (Figure 6A). However, we detected no correlation between IgG-targeting of the CTD2 or S1/S2 regions and severity (Figure 6C, 6E). In comparing the overall anti-spike levels with IgG-targeting of these regions, we detected a significant correlation between IgG-targeting of CTD2 and higher anti-spike IgG levels (Figure 6D), but did not observe any other significant correlation with the other regions (Figure 6B, 6F). However, targeting of CTD1 trended towards a significant positive correlation with overall anti-spike IgG levels (Figure 6B).

**Figure 6.**
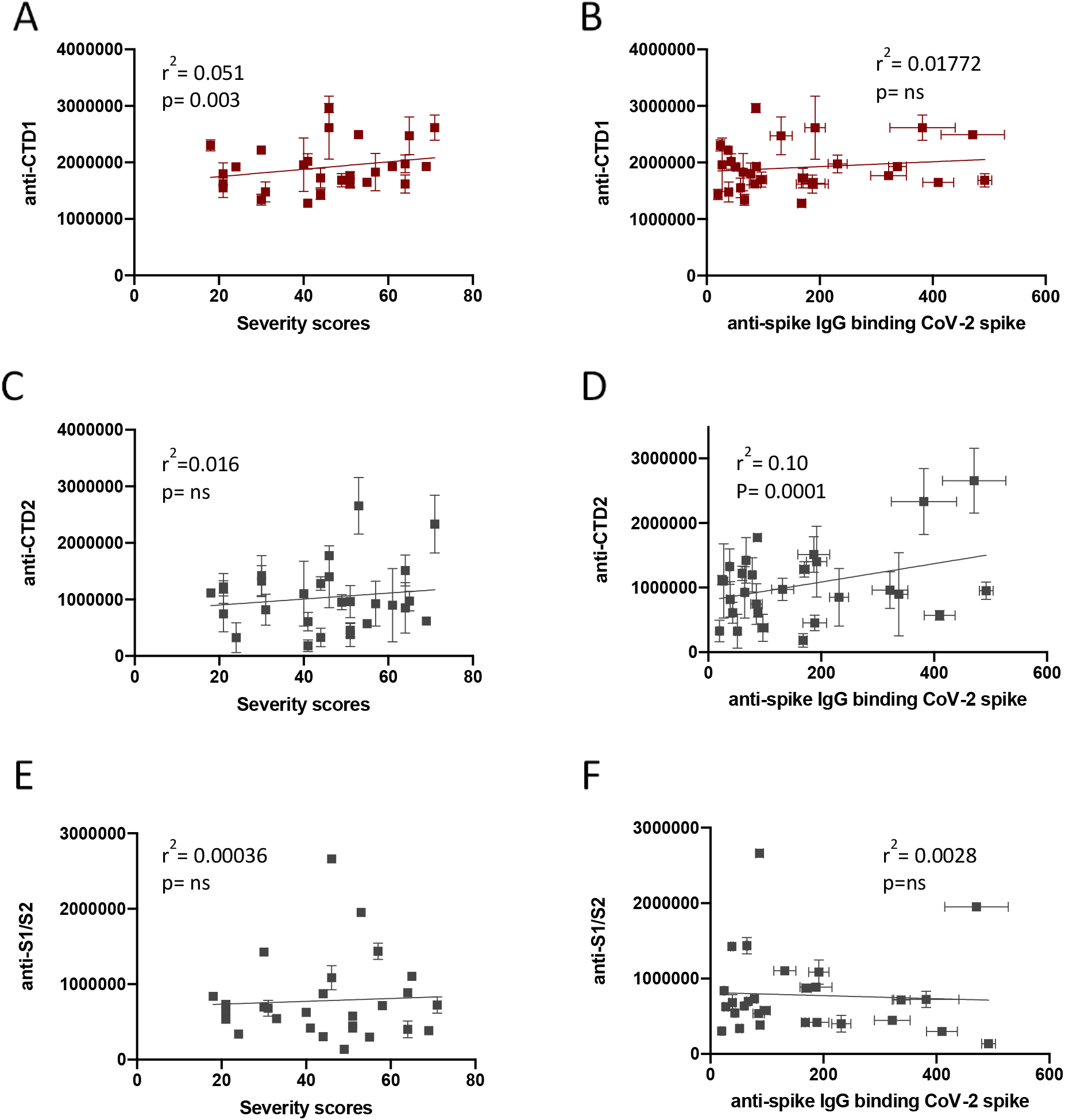
SARS-CoV-2 convalescent IgG differentially target non-conserved SARS-CoV-2 immunodominant epitopes. Graphs compare the levels of IgG binding to **(A**,**B)** CTD1, **(C**,**D)** CTD2, **(E**,**F)** S1/S2 regions versus total anti-spike levels, as quantified by cell-based binding assay. Graphs compare the levels of IgG-binding to **(A**,**B)** CTD1, **(C**,**D)** CTD2, **(E**,**F)** S1/S2 regions as detected by luminescent ELISA versus **(A, C, E)** severity scores or **(B, D, F)** total anti-spike levels quantified by cell-based binding assay. The levels of IgG targeting are the SEM of N=3 experiments are graphed to assess significance of slopes and r^2^ values.

### Multivariable analysis Identifies that severe non-hospitalized COVID-19 correlates with high FcγR activation and IgG targeting of spike protein’s S2’ fusion protein site

To gain insights into the associations between all of the Ab features tested and disease severity, we performed a Spearman’s chart correlation on all of the variables analyzed (Figure 7A). Our analysis recapitulated the significance of all of the previous findings in this study, such as the correlations between overall anti-spike levels, betacoronavirus cross-reactivity, and elevated levels of FcγR signaling. Additionally, this analysis allowed us to compare these variables with IgG-targeting of various immunodominant/functional regions.

**Figure 7.**
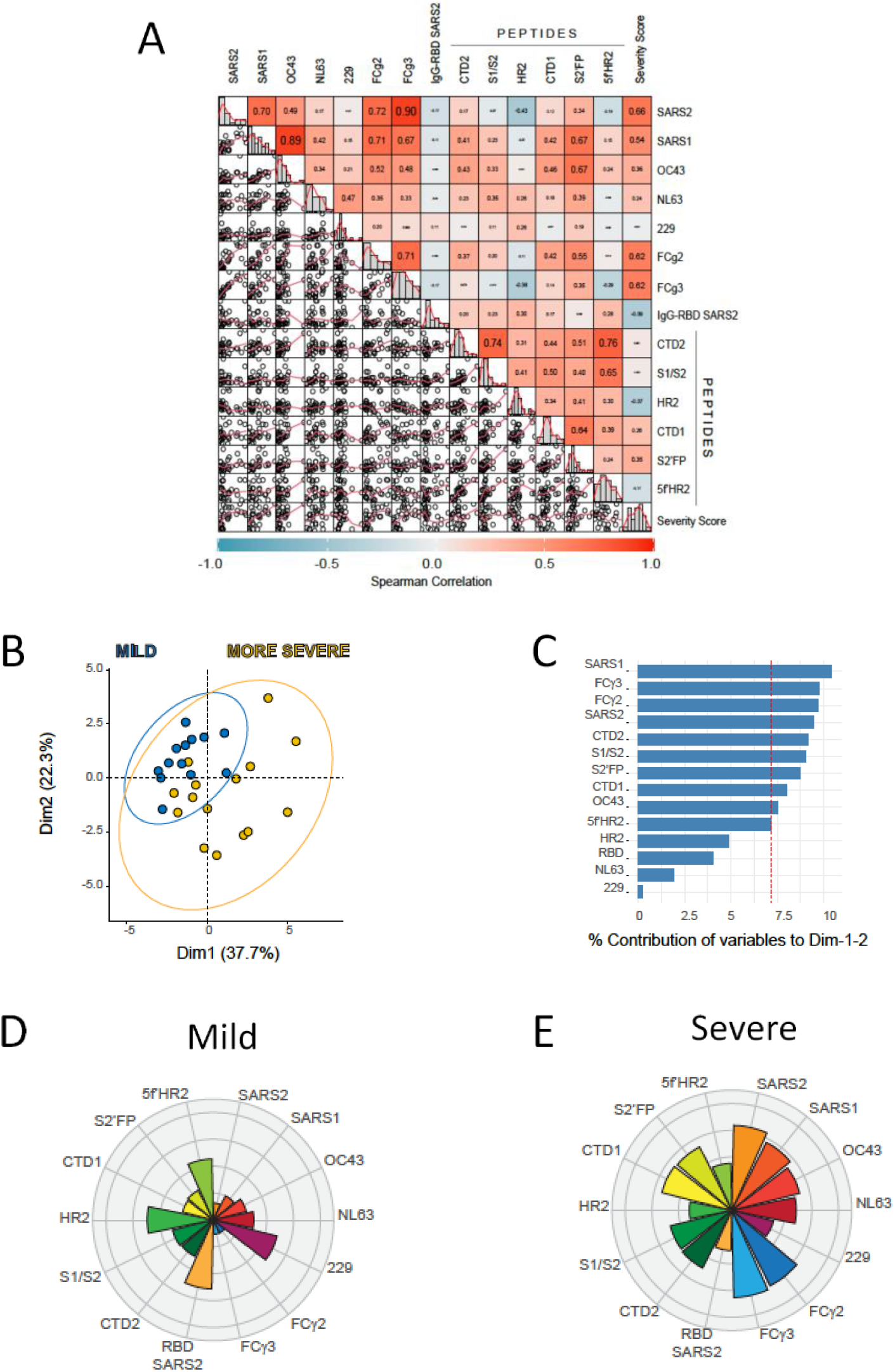
Multivariable analysis identifies that mild and more severe COVID19 is differentiated by distinct humoral immune profiles. **(A)** Scatter matrix chart summarizes the Spearman’s correlation (upper) and the scatter plots (lower) between all analyzed variables using the entire cohort (n=28). The Spearman’s r values are shown inside the colored squares and the scale of blue-to-red color indicates a negative-to-positive correlation. The small bar graphs (diagonal) represent the distribution of data for each variable **(B)** Biplot shows the principal component analysis (PCA) depicting the mild-scored (n=13) and more severe-scored (n=15) COVID-19 patients, according to their severity scores **(C)** The contribution of each variable to PCA for dimension 1 and 2 is represented by bars, and its threshold is indicated as a red dotted line. **(D**,**E)** Polar plots show the different profiles of humoral response for mild and more severe groups. Each bar in the plot represents the mean of z-scores for each variable.

In examining all convalescent donors, we observed that individuals with more severe COVID-19 were characterized by higher anti-SARS CoV-2 spike IgG (R=0.66), higher IgG cross-reactivity to betacoronaviruses (SARS1 R=0.54; OC43 R=0.36), and higher proinflammatory FcγR activation (R=0.62), along with higher levels of IgG targeting the CTD1 (R=0.26) and S2’FP region (R=0.35), as compared to mild donors (Figure 7A). Interestingly, higher severity scores were inversely correlated with the levels of IgG targeting the RBD and HR2 regions, suggesting that in severe individuals there was a relatively lack of IgG targeting neutralizing regions as a proportion of total anti-spike levels. Of note, the levels of betacoronavirus (OC43; SARS1) cross-reactivity were more closely associated with IgG targeting of SARS2 immunodominant regions as compared to the levels of alphacoronavirus (NL63; 229E) targeting (Figure 7A). Interestingly, while we observed that both FcγR2a and FcγR3a activation were positively correlated with betacoronavirus cross-reactivity, the levels of FcγR2a signaling were more highly correlated with IgG targeting the CTD1 (R2=0.42) and S2’FP (R2=0.55) regions, as compared to FcγR3a signaling (CTD1 R=0.14; S2’FP R=0.35). These findings suggest that seasonal hCoV IgG recall responses and epitope targeting contribute to the efficiency of FcγR signaling against the SARS-CoV-2 spike, especially in the case of FcγR2a activation.

To further explore the contribution of different IgG features, we performed an unsupervised analysis of principal components (PCA) using all 28 convalescent donors (Figure 7B, 7C). The results of this analysis showed that both Dimensions explain the 60% of variance and the severe individuals are clearly separated according to Dimension 1 (37.7% of variance), with a response focused on both FcγR2a and FcγR3a signaling, beta-coronaviruses anti-spike IgG (including SARS-CoV-2), and IgG-targeting of CTD1, CTD2, and S2’FP region. Interestingly, the individuals with more severe COVID-19 were highly represented on this Dimension, suggesting that cross-reactive IgG responses against betacoronaviruses—which included the induction of high Fcγ2a and Fcγ3a activation that targeted CTD1, CTD2 and S2’FP region—were/are predictive of severe disease. In contrast, mild individuals possessed different IgG responses, with lower levels on the features within Dimension 1, coupled with higher representation of Dimension 2 (22.3% of variance), which was primarily represented by IgG-targeting of the RBD, S1/S2 furin site, HR2 region, and 5f’HR2 regions. This analysis suggests a differential course of disease when the IgG response is directed against different regions of the SARS-CoV-2 spike.

Having identified several key IgG features segregating all 28 convalescent donor profiles based on severity, we next wanted to examine if mild and severe donor profiles were distinct and conducted a Spearman’s chart correlation of all Ab variables using only mild or severe donors (Supplementary Figure 4A, 4C). In comparing the profiles of mild and severe individuals, the most striking difference was the lack of a correlation between FcγR2a activation and IgG targeting of immunodominant regions in mild individuals. This was in sharp contrast to the profiles of severe individuals, where FcγR2a activation was highly correlated with targeting of CTD1, CTD2, and S2’FP regions (Supplementary Figure 4C).

To better identify any correlations with anti-spike, anti-RBD, and immunodominant epitope targeting amongst mild and severe profiles, we next conducted a separate Spearman’s chart correlation on only the anti-spike region IgG targeting profiles. We observed that individuals with milder infection possessed anti-spike IgG levels that were inversely correlated with the levels of S2’FP (R=-0.27) and CTD1 (R=-0.41) region IgG targeting (Supplementary Figure 4B). Interestingly, regarding RBD-targeting in mild donors, we did not observe a relationship between anti-RBD levels and HR2 region targeting (R=-0.077), possibly suggesting two divergent protective signatures, one recall and one *de novo*, respectively. In severe profiles, we observed that total anti-spike IgG levels were inversely correlated with IgG targeting the S1/S2 (R=-0.39) and HR2 region (R=-0.42) and positively correlated with S2’FP (R=0.37) region IgG-targeting (Supplementary figure 4D). Interestingly, anti-RBD IgG levels in severe donors were not strongly correlated with the levels of IgG-targeting of any immunodominant regions.

To further elucidate the overall profiles between mild and severe individuals, the mean of Z-score values for each antibody feature was represented on a polar plot graph (Figure 7D, 7E). Notably, for mild individuals, the IgG recognition of betacoronaviruses was lower than in severe individuals (Figure 7D, 7E). However, in comparing the relative targeting of betacoronaviruses vs alphacoronaviruses, we observed that mild individuals expressed higher Z scores for IgG targeting alphacoronaviruses, with 229E being more highly targeted as compared to NL63 (Figure 7D). In examining only severe profiles, the inverse pattern was observed, where more severe individuals possessed higher Z scores for betacoronavirus-targeting when compared to alphacoronaviruses (Figure 7E). In support of the PCA analysis relating disease severity to FcγR function, the polar plot shows that individuals with mild disease had lower FcγR2a and FcγR3a enrichment scores (Figure 7D). Crucially, despite severe donors possessing higher overall levels of anti-spike IgG, the enrichment scores for RBD IgG-targeting was higher in mild donors (Figure 7D). This observation may imply that the elevated levels of anti-spike Abs observed in more severe individuals could be attributable to inefficient non-neutralizing responses against SARS-CoV-2, since milder profiles are seen to be enriched for a higher proportion of IgG targeting the RBD, HR2, and HR2 flanking regions, which have been previously shown to neutralize both SARS1 and SARS2 [55-57, 78-83]. Additionally, Z-scores in mild individuals indicate a relative lower enrichment of IgG targeting the CTD1 and S2’FP regions, both of which were significantly correlated with elevated FcγR activity in severe individuals (Figure 7D). In severe individuals, the inverse was observed: IgG targeting CTD1 and S2’FP regions was enriched, and HR2 region and 5’fHR2 targeting was decreased, relative to total anti-spike IgG (Figure 7E). Taken together, these data suggest that efficacy of IgG responses—and corresponding disease severity—is highly dependent on the specific epitopes targeted in the SARS-CoV-2 spike protein, which in turn may be influenced by prior hCoV exposure.

## Discussion

COVID-19 is expected to remain an ongoing global threat, driven in part by emerging variants and challenges to vaccine distribution worldwide. Unlike many other pathogens that induce long-lived immunity, seasonal coronaviruses have been observed to induce transient immunity, which rapidly wanes following natural infection. Vulnerability to seasonal coronavirus reinfection is the norm starting 6 months to 3 years after each seasonal hCoV infection [84]. Evidence has emerged that SARS-CoV-2 reinfection is possible after 3-6 months, particularly with emerging variants of concern [85-90]. Moreover, vaccine-induced immunity wanes significantly after 6-8 months [91, 92]. This uncertainty surrounding the duration of immunity (whether natural or vaccine-induced), combined with the continuing evolution of novel SARS-CoV-2 mutants, reinforces the need to continue to identify the immune correlates of protection in SARS-CoV-2 infections in humans. Identifying the protective correlates of Ab-targeting is particularly important, as they may provide new insights into mechanisms of protection that can be leveraged for more effective or broadly-acting therapeutics and vaccines.

Early in the pandemic, researchers noted an unusual aspect of COVID-19 disease course; namely, some patients were showing IgG within days of contracting SARS-CoV-2. [7, 9, 11] Furthermore, early anti-spike IgG was associated with disease severity—not protection. Higher anti-spike IgG titers, along with elevated levels of proinflammatory cytokines, correlated with more severe disease in hospitalized individuals [9, 12]. In this study, we observed that increasing anti-spike IgG titers were correlated with higher symptom severity scores, substantiating and extending this phenotype to non-hospitalized individuals. We detected significantly higher levels of anti-spike IgG using both ELISA and cell-based assays. Of note, the cell-based assay detected anti-spike IgG with greater sensitivity than ELISA. Other studies have also observed higher sensitivity with cell-based detection assays [93, 94]. This may be attributable to the display of conformational epitopes that are dependent on tertiary or quaternary structure which occur in natively folded trimeric spike protein. Moreover, commercial recombinant antigens produced in bacteria or insect cells may not accurately reflect the glycosylation pattern produced in human cells [95, 96]. This difference in sensitivity should be considered when selecting IgG assays for varying research and clinical uses.

In this study, we observed a significant positive correlation between severity, anti-SARS-CoV-2 spike IgG levels, FcγR signaling, and IgG cross-reactivity with other betacoronaviruses; with the highest levels of FcγR-signaling and IgG cross-reactivity in the donors with the highest symptom severity scores. These findings suggest that anti-spike IgG may be contributing to COVID-19 severity through the amplification of inefficient and cross-reactive IgG responses against seasonal hCoVs, potentially elevating proinflammatory cytokine levels through IgG-induced FcγR activation. Early in the pandemic, some researchers had optimistically posited that Abs from previous hCoVs might be protective against SARS-CoV-2. However, subsequent studies identified that pre-pandemic and SARS-CoV-2 naïve samples did not mediate effective virus neutralization, and predominantly targeted the S2 subunit of the SARS-CoV-2 spike protein [58-60]. Importantly, the RBD region, located in the S1 subunit of the spike protein, interacts with the human ACE2 receptor to mediate viral attachment and entry into host cells [74]. In studies characterizing neutralizing Abs from convalescent donors, Abs targeting the SARS-CoV-2 RBD region were shown to potently inhibit virus replication and control infection *in vivo* [55-57]. In contrast to the S1 subunit that is largely non-homologous with seasonal hCoVs, the S2 subunit contains the highest sequence and structural similarity with seasonal coronaviruses [54]. Therefore, the seroreactivity observed in some pre-pandemic and naïve donors is likely a result of SARS-CoV-2 cross-reactive humoral memory responses against seasonal hCoVs. In light of this possibility, we initially hypothesized that any seasonal hCoV recall responses amplified during acute infections would favor S2 subunit targeting of the spike protein. Moreover, if these recall Abs dominated the humoral response within an individual, we further hypothesized this would lead to an inefficient non-neutralizing response (ala Original Antigenic Sin), correlating with worse outcomes. In line with this hypothesis, the early and elevated levels of anti-spike IgG observed in severe COVID-19 patients could be a consequence of amplifying cross-reactive recall responses against S2 targeting non-neutralizing epitopes. In this manner, anti-spike Ab titers may increase due to inefficient Ab-targeting and uncontrolled virus replication, leading to an elevated inflammatory environment and worse outcomes. Whereas *de novo* responses (i.e., S1/RBD/novel sequence)—which are broader and inherently SARS-CoV-2 sequence-specific—would be protective.

Despite these reasonable assumptions, in screening the levels of IgG targeting highly conserved immunodominant regions, we detected no uniform association between high sequence identity and severity. We instead observed that IgG-targeting of two highly conserved regions (HR2 region, 5’HR2 flanking region) was significantly correlated with milder infections, while targeting of another conserved region (S2’FP site) was correlated with more severe infections. Furthermore, several studies have reported that both HR2 and S2’FP are among the most immunodominant regions in the spike protein [62, 68]. Interestingly, SARS-CoV-2 naïve individuals have been shown to possess Abs that react to both of these regions [68], demonstrating that prior exposure to homologous coronaviruses could give rise to cross-reactive memory responses. Notably, (IgG/Ab) targeting of HR2 region and 5’HR2 flanking region was previously shown by multiple studies to mediate neutralization of both SARS1 and SARS2 *in vitro* [78-83]. Moreover, peptide-based SARS-CoV-2 fusion inhibitors targeting the HR1-HR2 fusion complex within regions that are highly conserved with OC43 have been shown to potently inhibit virus replication [97]. However, only one study reported that Abs targeting the S2’FP region neutralize SARS-CoV-2 *in vitro* [98]. While the neutralizing activity associated with IgG-targeting of the HR2 region is consistent with our data showing that IgG-targeting of the HR2 region is correlated with milder infections, IgG-targeting of the S2’FP region correlated with more severe infections. Importantly, studies identifying the HR2 and S2’FP regions as immunodominant did not examine the relationship between immunodominance and infection severity [68]. Therefore, despite the potential for Ab-targeting of the S2’FP region to mediate some level of neutralization against SARS-CoV-2, whether or not targeting this region correlates with milder outcomes and presumably more effective control of infection *in vivo*—remains undefined.

While we did not examine IgG neutralization activity in this study, we did examine the capacity of IgG to induce FcγR activation, a prerequisite for inducing non-neutralizing effector functions like proinflammatory cytokine release. In this analysis, we identify that increased FcγR-signaling is highly correlated with higher IgG targeting of the S2’FP region and more severe COVID-19. This finding provides a rationale as to why targeting this epitope may correlate with severity, since FcγR activation in the context of ADE can induce proinflammatory cytokines, which are highly associated with severe hospitalized cases of COVID-19. Notably, anti-SARS-CoV-2 Ab-targeting of the S2’FP region was shown to be one of the most immunodominant regions targeted within a severe COVID-19 patient after prolonged hospitalization and intensive care [61]. Altogether, our data shows clear correlations between all three of the immunodominant conserved regions and severity, whether inverse or positive. In contrast, we only noted a significant correlation between IgG-targeting and severity with one of three non-homologous regions.

Importantly, these data demonstrate that the contribution of recall antibodies to COVID-19 disease severity is nuanced; IgG-targeting of some conserved neutralizing epitopes may be protective, while targeting of other conserved epitopes may activate high levels of FcγR-mediated proinflammatory signaling, which in some instances may be detrimental. Importantly, while cross-reactive responses, like those against the S2’FP region, may contribute to the severity of infection, cross-reactivity in and of itself may not be the sole factor in governing inefficient humoral responses during SARS-CoV-2 infections. These findings highlight that the relationship between seasonal hCoV humoral cross-reactivity and COVID-19 disease outcomes may be highly dependent on an individual’s pre-existing hCoV IgG repertoire prior to becoming infected with SARS-CoV-2. Therefore, further examination of how specific IgG-targeting and pre-existing memory responses influence and contribute to pathogenesis *in vivo* will be key to fully understand how cross-reactivity contributes to effective humoral responses.

Beyond defining the spike protein immunodominant regions based on their similarity to betacoronaviruses (OC43), these regions also represent known functional regions within the spike protein. Examining the profiles of IgG-targeting among convalescent individuals in the context of functional regions, we observed that IgG-targeting of RBD, HR2, and HR flanking regions significantly correlated with lower severity scores and—in the case of HR2 region-targeting—also correlated with lower levels of anti-spike IgG. This latter finding suggests that humoral responses with a higher proportion of IgG-targeting against the HR2 region may be highly efficient at controlling infection, since lower anti-spike titers were independently associated with milder infections in this study and others [7-9, 12]. In previous studies examining Ab-targeting of the RBD and HR2 regions, targeting both of these regions have been shown to mediate virus neutralization [55-57, 83]. In experiments conducted with SARS1, targeting of the HR2 flanking region also neutralizes virus replication [79, 82]. Interestingly, when we examined the levels of anti-RBD IgG targeting in mild donors, we observed a positive correlation with S1/S2 furin site and HR2 flanking region IgG-targeting, but not with HR2 region targeting. Therefore, it is possible that targeting the HR2 flanking region sufficiently disrupts the interaction between HR2 and HR1, thereby preventing viral fusion. Moreover, when we compared the levels of RBD vs immunodominant epitope targeting in mild donors, we observed no correlation between RBD and HR2 region IgG-targeting, despite both being correlated with mild disease. This raises the possibility that one profile of neutralization predominantly arises in some individuals and not in others, possibly influenced by whether or not humoral responses are predominately recall or *de novo*. A closer examination of the relationship between RBD and HR2 region neutralization will be the subject of future studies.

In regard to which IgG-targeting profiles correlated with overall severity amongst all convalescent donors, we detected a correlation with IgG targeting the non-conserved CTD2 region and the conserved S2’FP region. In further examining the profiles of only severe donors, we resolved that these profiles were characterized by higher levels of IgG targeting S2’FP region, with an absence of IgG targeting the S1/S2 furin site and HR2 region (Supplementary Figure 4C). Notably, milder IgG profiles were not only characterized by higher levels of RBD and HR2 region IgG-targeting, but also lower proportions of CTD1 and S2’FP region targeting in relation to overall anti-spike titers (Figure supplementary 4B). These findings strongly suggest that targeting of the S2’FP region—coupled with the absence of anti-RBD and/or anti-HR2 region IgG-targeting—is linked with severity; further implying that these IgG profiles represent inefficient responses against the SARS-CoV-2 spike.

Altogether, these findings suggest that humoral memory responses contribute to COVID-19 disease severity, conferring either protection or risk depending on epitope targeting. This may explain the atypical bimodality of COVID-19 disease severity—an observation which was previously obscured by aggregate data/epitope analysis. The correlation we observed with IgG-targeting of the spike protein highlights an important point since to date it’s been unclear why some people respond so severely to infection while others have a mild infection. Thus far, it has been challenging to enumerate the underlying factors that accurately predict disease outcome, even among individuals who are at high risk of severe symptoms. The findings reported here have the potential to help improve prediction accuracy and explain the underlying mechanisms of COVID-19 pathogenesis, as pre-existing cross-reactive immunity with seasonal coronaviruses is correlated with, and likely contributes to, disease outcome. This epitope-based Original Antigenic Sin immunosurveillance, or OASiS, could be readily adapted to a clinical prognostic, offering a novel approach for the prediction of disease severity risk, as well as vaccine efficacy, on a patient-by-patient basis. Dissecting the interplay between these epitopes and the efficiency of IgG-mediated immunity will be the focus of future work.

Beyond defining how IgG-targeting of immunodominant regions correlates with COVID-19 outcomes, we also show that higher levels of FcγR2a and FcγR3a activation correlate with more severe SARS-CoV-2 infections. Interestingly, both FcγR2a and FcγR3a are correlated with more severe dengue pathogenesis in humans and animal models, and have been shown to promote ADE effects *in vitro* [33-35, 99]. In SARS1 infections, *in vitro* experiments using IgG from severe SARS1 patients have been shown to induce the FcγR-mediated activation of proinflammatory cytokines from macrophages [13]. In studies examining the role IgG in severe COVID-19, higher levels of the afucosylated glycoform of IgG—which inherently possess a higher capacity to activate FcγRs—have been correlated with more severe COVID-19 [22, 24, 100], and have also been shown to activate cytokine expression from macrophages [24, 100]. Notably, the levels of afucosylated anti-spike IgG were correlated with higher C-reactive protein and IL-6 levels in severe hospitalized COVID-19 patients, as compared to convalescent non-hospitalized individuals with milder symptoms [22]. Importantly, afucosylated anti-spike IgG were also shown to possess higher affinity for FcγR3a and induce high levels of IL-6 and TNF-α release from monocytes and macrophages *in vitro* [24, 100]. These findings suggest that FcγR-activation contributes to the proinflammatory profile in COVID-19 patients. However, these results do not define the contribution of FcγR effector functions in the control of SARS-CoV-2 infections *in vivo*. Whether or not this increased Fc-activation is a compensatory mechanism for controlling virus replication in the face of inefficient IgG neutralization, or the driver of severe pathogenesis though increasing inflammation via the induction of proinflammatory cytokines, remains an open question.

Interestingly, in murine models of SARS-CoV-2 infection, the passive infusion of neutralizing IgG directed against the RBD region was shown to effectively protect animals when administered prophylactically, regardless of whether or not the Fc-binding activity of the neutralizing IgG remained intact [101]. However, in therapeutic intervention models (i.e., post-exposure), the ablation of Fc-activity from neutralizing IgG significantly decreased the ability of IgG to effectively control infection *in vivo* [101]. Similar findings have been observed with other pathogens, such as Ebola, HIV, and influenza. In these studies, the therapeutic efficacy of neutralizing IgG was also shown to depend on intact FcγR activity [42-45, 102]. Moreover, in our own previous studies with hantaviruses, we identified an IgG Ab, JL16, which possessed highly efficient FcγR activation and neutralizing activity. When tested in animal models of hantavirus infection, the passive administration of JL16 as a therapeutic during acute infection yielded more effective control of virus replication when compared against an antibody with more potent neutralization [72]. Altogether, these studies suggest that in the context of active viral replication and established infection, IgG FcγR activity may be required for potent control of viral infections *in vivo*.

In this study, we observed that more severe SARS-CoV-2 infections were characterized by lower levels of IgG targeting the RBD and HR regions. This suggests that high FcγR activation in the absence of high titers of neutralizing IgG may be deleterious. One caveat to the previous studies that found FcγR activity to be critical for effective humoral immunity is that all of those studies were conducted with neutralizing IgG. Therefore, we cannot rule out that FcγR activation in the absence of a neutralizing response does not contribute to severe COVID-19 pathogenesis, as may be the case with individuals with anti-S2’FP dominant IgG profiles. Therefore, it will be important to dissect how IgG-targeting of neutralizing and non-neutralizing epitopes influences the relationship between FcγR activation and disease outcomes. Additionally, it will be important to examine the contribution of FcγR activation to systemic inflammation and control of virus replication in animal models that faithfully mimic human FcγR-signaling effects.

In conclusion, the data presented here demonstrate that immunological history—particularly antibody repertoire from seasonal betacoronaviruses—predicts and potentially determines COVID-19 disease severity. Specifically, an anti-HR2-dominant Ab profile represents an efficient protective recall response, an anti-S2’FP dominant Ab profile suggests a deleterious inefficient recall response, and a predominantly anti-RBD profile likely reflects a protective *de novo* response. These data become particularly important in assessing epitope-targeting early in disease, which could allow earlier interventions with treatments like monoclonal Ab therapies, preventing progression to ARDS. Additionally, assessing the humoral profiles induced by vaccination will be important. Vaccination will boost SARS-CoV-2-specific IgG, both novel and recall responses, initially giving some level of protection, but could lead to higher frequencies of breakthrough infections over time if inefficient IgG-targeting remains the predominant response. Fortunately, vaccine studies have shown that current mRNA vaccines appear to induce high titers of RBD-specific antibodies; however, anti-RBD-targeting Abs are predominantly induced only after the second dose [103, 104]. This finding suggests potentially meaningful variation in IgG profiles induced by different vaccines, particularly multi-vs single-dose regimens, such as J&J. Therefore, it will be important to assess the IgG-targeting profiles induced by all available vaccines, as this type of analysis may help to explain why some vaccines induce effective anti-SARS-CoV-2 Abs while others do not produce lasting or robust protection, such as SinoVac. Additionally, it will be important to examine the ratios of RBD to other immunodominant epitope IgG targeting and assess the longevity of immunity and the frequencies of breakthrough infections over time. Altogether, epitope-profiling may be essential to ensure long-term vaccine-induced protection in individuals, as individuals with inefficient IgG profiles (i.e., anti-S2’FP-dominant), even after vaccination, may require an RBD-specific or HR2-specific vaccine to change their targeting profiles and induce long-lived protective humoral responses. Promisingly, HR2 offers a good target for universal hCoV treatment and vaccine design.

## Materials and Methods

### Subjects and samples

Demographic data (age, gender, COVID-19 RT-PCR status), and collection site and date of the serum/plasma donors studied herein are described in Table 1. COVID-19 convalescent donor samples (n = 28) and COVID-19 negative donor (n=20) were collected at Ichor Biologics facility in New York City, USA. SARS-CoV-2 negative donors had no history of positive SARS-CoV-2 PCR test or serology test, and had not experienced any symptoms of infection in at least the 5 months prior to blood collection. Blood samples were collected after obtaining signed informed consent in accordance with institutionally approved IRB protocols. Convalescent COVID-19 patient samples were collected from donors 2-10 weeks after the onset of symptoms. A history of COVID-19 symptoms and symptom intensities, along with current medications and history of pre-existing conditions, were collected through participant questionnaires completed at the time of blood draw. Lithium heparin-coated tubes were used for blood collection and plasma was isolated using Ficoll-Hypaque (GE Healthcare; 17-1440-03) in accordance with manufacturer’s instructions. Polyclonal IgG was isolated from 200μl of donor plasma using a protein A/G spin column kit, followed by desalting using Zeba spin columns according to manufacturer’s instructions (ThermoFisher Scientific; 89892). IgG yields were quantified using an Easy-Titer Human IgG Assay Kit (ThermoFisher Scientific; 23310). Remaining deidentified plasma samples were aliquoted and stored at -80C.

### SARS-CoV-2 Spike-ELISA

High binding capacity 96-well plates (Nunc) were washed and coated with 50 μl per well of 2μg/ml of recombinant spike protein (Sino Biological; 40589-V08B1-B), diluted in 0.1% BSA, 0.05% Tween20 TBST ELISA wash buffer (ThermoFisher Scientific; N503). Plates were coated for 2 hours at room temperature while shaking at 500 rpm on a Benchmark Orbishaker™. Plates were then washed twice with ELISA wash buffer to remove any excess unbound spike protein and blocked with 2% BSA (ThermoFisher Scientific; 37525) in ELISA wash buffer overnight at 4°C. After overnight blocking step, plates were washed twice and incubated with 5μg/ml of donor-derived polyclonal IgG for 1 hour at room temperature. After incubation, plates were washed three times and incubated for 30 minutes at room temperature with cross-absorbed goat anti-human IgG-horseradish peroxidase (HRP)-conjugated secondary antibody (ThermoFisher Scientific; A18811) diluted to a 1:2500 dilution in ELISA wash buffer. After being washed again twice, 100μl of TMB substrate solution was added to each well for 15mins and then 100μl of 0.18M H_2_SO_4_ (ThermoFisher Scientific; N600) was added to stop the reaction. The optical density at 450 nm (OD450) was measured using a BioTek Powerwave HT plate reader using Gen5 software. Assay background was established using anti-human secondary Ab alone without donor IgG, which was subtracted from OD values of all samples tested.

### Anti-spike protein IgG determination using a cell-based assay

To quantify the levels of IgG binding various coronavirus spike proteins, 293T cells were transfected with SARS-CoV-2 (Sino Biological; VG40589-CF)(Genbank YP_009724390.1), SARS-CoV-1 (Sino Biological; VG40150-CF)(Genbank AAP13567.1), OC43 (Sino Biological; VG40607-CF)(Genbank AVR40344.1), NL63 (Sino Biological; VG40604-CF)(Genbank APF29071.1), or 229E (Sino Biological; VG40605-CF)(Genbank APT69883.1) spike protein expression vectors. For this assay, 2×10^6^ 293T cells were plated in 10cm plates and incubated at 37°C overnight. The next day, 4μg of coronavirus spike expression vectors were transfected into 293T cells using Polyjet™ transfection reagent (SignaGen; SL100688) according to manufacturer’s instructions. After 48 hours, 1×10^5^ 293T cells were plated per well into round bottom 96-well plates. Cells were then washed and incubated with 10μg/ml of convalescent donor-derived IgG or negative donor control IgG and incubated at 4°C for 45 minutes. After primary Ab incubation, IgG opsonized cells were washed and incubated with 3μg/ml of an APC-conjugated anti-human total IgG secondary Ab (Invitrogen, catalog A21445) at 4°C for 25mins. Cell were then washed again with PBS and LIVE/DEAD™ Fixable Violet Stain (Invitrogen; L34964A) was used to stain cells for 10 mins in the dark at RT. Lastly, cells were washed twice and fixed with 1.0% paraformaldehyde in PBS and analyzed by flow cytometry (BD LSRFortessa X-20). The data were quantified using Flow Jo software (Tree Star, Inc). The IgG-binding index was calculated by multiplying the percentage of anti-spike IgG positive cells by the Median fluorescent intensity (MFI) of APC signal, as normalized to the average MFI of negative control IgG. To ensure that the relative differences between patient-derived IgG were maintained, all IgG were tested in parallel on the same day for each replicate.

### Fc-gamma receptor signaling assay

FcγR2a and FcγR3a signaling was assessed using a reporter cell co-culture system that we have previous used to assess FcγR signaling in response to viral antigens ([70, 71]). For this assay, 293T cells are transfected with SARS-CoV-2 spike expression vector and co-cultured with either a FcγR2a, or FcγR3a, CD4^+^ Jurkat reporter cell line, which expresses firefly luciferase upon FcγR activation. For this assay, 1×10^5^ SARS-CoV-2 spike-expressing 293T cells were plated in each well of a 96-well round bottom plate. The cells were then preincubated with a 5-fold dilution series of convalescent donor-derived IgG starting at a maximum concentration of 25μg/ml. IgG opsonized 293T cells were then co-cultured with FcγR2a or FcγR3a reporter cells at a 2:1 reporter-to-target cell ratio for 24hrs at 37°C. After 24 hours, all cells were lysed with cell lysis buffer (Promega; E1531), and the levels of firefly luciferase activity determined using a luciferase assay kit according to manufacturer’s instructions (Promega; E1500). To quantify background (i.e., IgG activation-independent) luciferase production, reporter cells were co-cultured with the spike-expressing 293T cells in the absence of any IgG. Background levels were subsequently subtracted from the signal to yield IgG-specific activation in relative light units (RLUs). Luminescence was measured on a Cytation 3 image reader using Gen5 software.

### Immunodominant Epitope IgG-binding assay

For this assay, N-terminus biotinylated peptides were synthesized by Genscript. N-terminal GSGS linker sequence was added to all peptide sequences. The RBD peptide contained a C-terminal avitag (GLNDIFEAQKIEWHE), for biotinylation via BirA enzyme; a Protein C tag (EDQVDPRLIDGK), and a polyhisidine tag (HHHHHHHHHH), to enable immobilized metal affinity chromatography purification. Lyophilized peptides and RBD were initially resuspended in DMSO and then used to make 5μg/ml working dilutions in TBST ELISA wash buffer. Pierce™ white streptavidin-coated high binding capacity 96-well binding plates (ThermoFisher Scientific; 15502) were washed twice with ELISA wash buffer and coated with 5μg/ml of biotiylated peptides at room temp. Plates were coated for 2hrs while shaking at 500rpm on a Benchmark Orbi-Shaker™. After incubation, plates were washed three times and blocked with 2% BSA blocking solution diluted in wash buffer and incubated at 4°C overnight. After incubation, plates were washed three times and incubated with 5μg/ml of donor-derived IgG for 1 hour at room temp. After primary IgG incubation, plates were washed 3 times and incubated for 25mins with 100μl of Invitrogen™ cross-absorbed, F(ab’)_2_, goat anti-human IgG secondary Ab (ThermoFisher Scientific; A24470) diluted 1:2500 in ELISA wash buffer. Plates were then washed again three times and developed using a SuperSignal™ ELISA Pico Chemiluminescent Substrate (ThermoFisher; Cat#37069) and the level of luminescence detected using a Cytation 3 image reader luminometer using Gen5 software.

### Statistical analysis

Statistical and data analyses were performed using GraphPad Prism 8.4.3, R 4.0.4, and R Studio 1.4.1103. Graphs were generated in Prism and R Studio and statistical differences between two groups were calculated by Mann-Whitney U-test. Statistical significance was defined as * p < 0.05; ** p < 0.01; *** p < 0.001, and **** p < 0.0001. Scatter plots, bar graphs, heatmaps, and polar plots were visualized with ggplot2 (v3.3.3 R Studio). Correlation analysis (in Figure 7 and S4) were performed using the R package “correlation” (v0.6.0) in R Studio. Polar plots represent the value of different variables normalized to the Z-score of data. Each variable was mean-centered and then divided by the standard deviation of the variable to ensure each variable had zero mean and unit standard deviation. Unsupervised principal components analysis (PCA) was performed in R. The completed data were scaled to unit variance using FactoMineR (v2.4 R studio). The PCA results were extracted and visualized using factoextra (v1.0.7 R Studio). Outlier exclusion was performed using Prism.

## Author Contributions

Study design was done by JLG, RAA, RAB, CDB, MC, FF, MIB.

Experimental work was performed by JLG, FB, SM, JWB, RAA.

Data analysis was performed by JLG, MM, SM, CDB, RAB, RAA.

Reagent development was done by JLG, RAA, JWB, and CDB.

Donor plasma acquisition was managed by SM, JLG, & RAA.

The manuscript was written by JLG, RAA, RAB and the final version was approved by all authors.

## Acknowledgments

We would like to thank Sheila O’Donoghue RN of the New York Presbyterian Hospital and Beth Sferrazza RN of Northwell Health/LIJ for their help in coordinating and conducting blood draws for this study. We would like to thank Drs. Svenja Weiss of the Icahn School of Medicine at Mount Sinai and Biliana Lozanoska-Ochser of the Sapienza University of Rome for their critical review of this manuscript. We would like to thank Dr. Server Ertem of the Jacobs Cornell-Technion Institute and Katena Oncology for his technical and scientific insight on diagnostic design and Dean Greg Morrisett PhD of Cornell Tech for his generous institutional support.

