## Supplementary Figures for "Humoral immune responses against seasonal coronaviruses predict efficiency of SARS-CoV-2 spike targeting, FcγR activation, and corresponding COVID-19 disease severity"

### Supplemental Figures

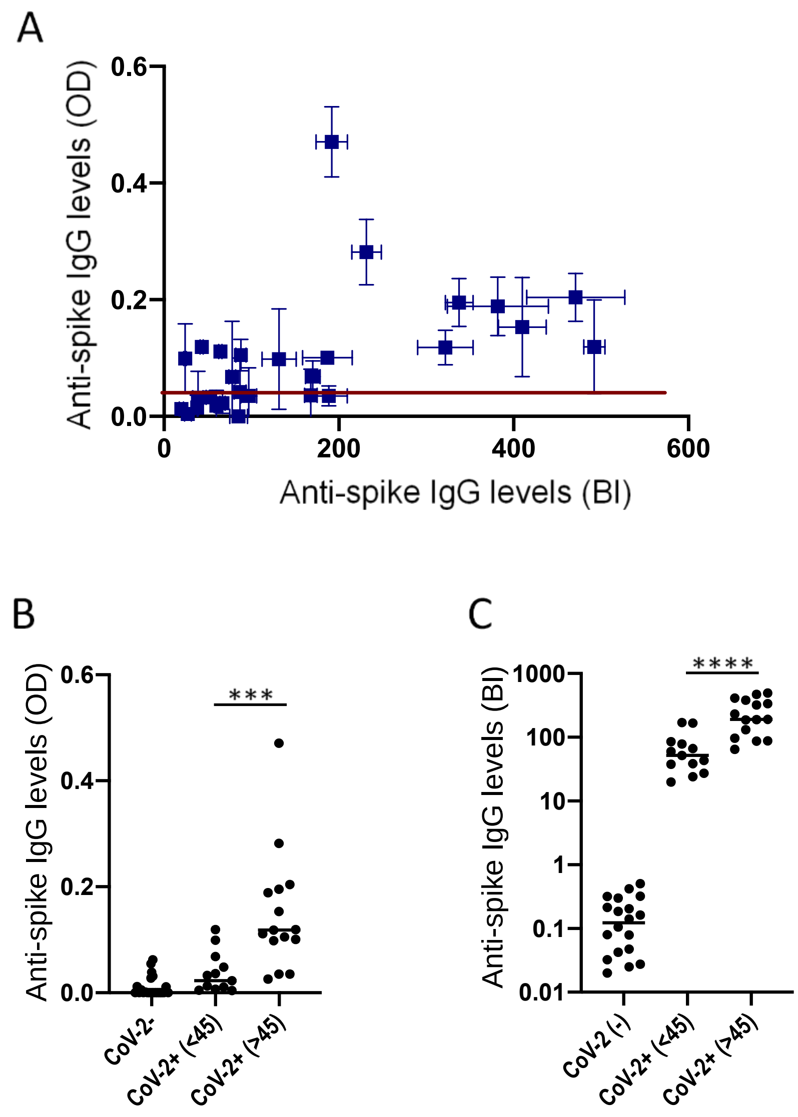


**Supplementary Figure 1. Comparison of anti-spike IgG levels in convalescent donors as quantified by ELISA vs Cell-based assays.**
**(A)** Anti-spike IgG titers in SARS-CoV-2 naïve (CoV-2-) and SARS-CoV-2 positive convalescent (CoV-2+) donors, as quantified by recombinant spike protein ELISA vs Cell-based binding assay. Red line represents 3-fold above the mean anti-spike levels of naïve (CoV-2-) donors, as quantified by ELISA. **(B&C)** Levels of anti-spike IgG titers as quantified by (B) ELISA, or (C) Cell based IgG binding assay. SARS CoV-2 naïve (CoV-2-) and convalescent (CoV-2+) donors are shown with convalescent donors split by COVID19 severity scores. The SEM of N=3 experiments are shown.

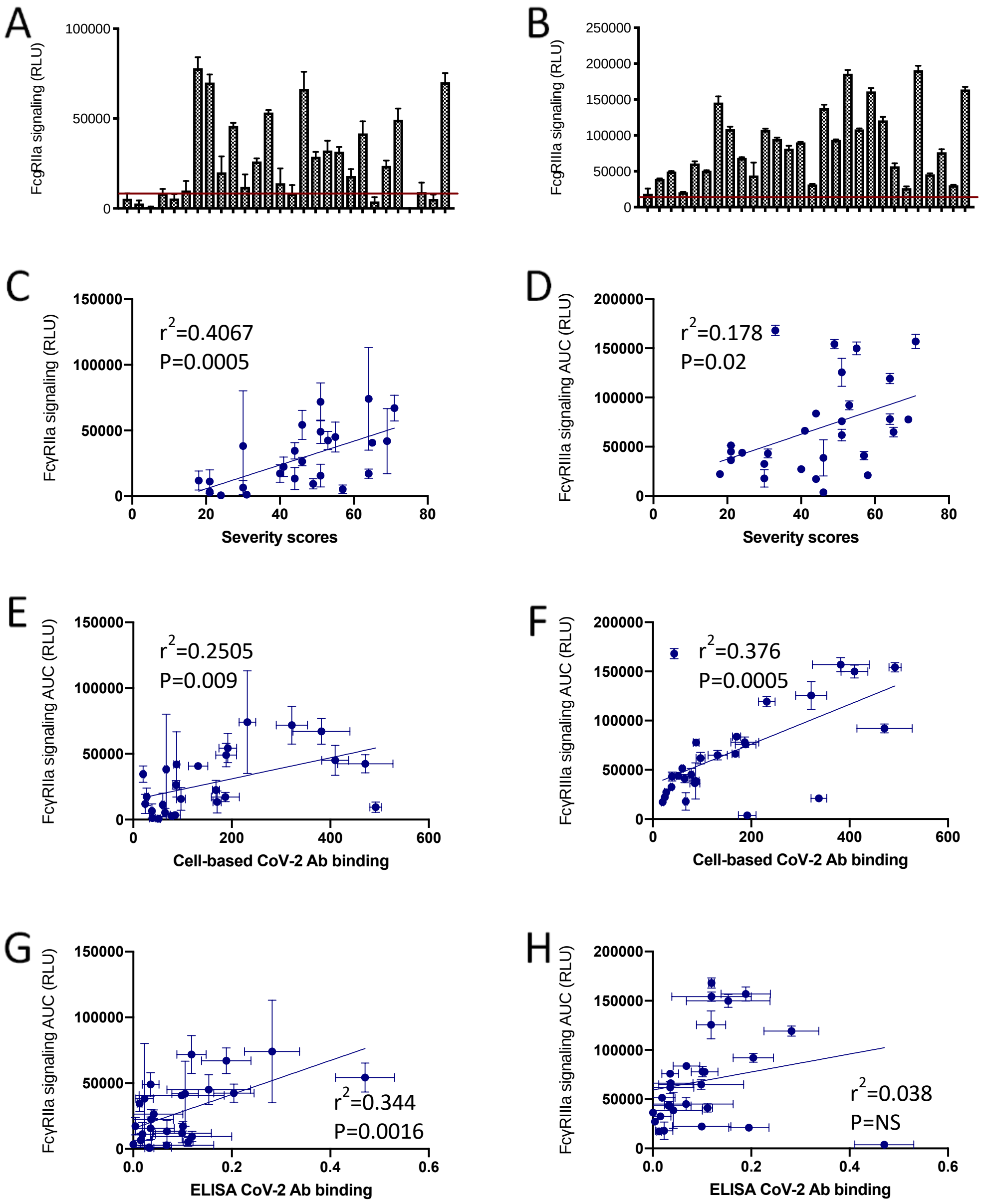


**Supplementary Figure 2. Levels of anti-spike IgG-induced FcγR-activation correlates with COVID-19 severity and anti-spike titers.**

Figure depicts the levels of FcγR-signaling induce by purified IgG derived from SARS-CoV-2 convalescent donors in response to SARS-CoV-2 spike protein expressed on the surface of 293T cells. Graphs show the levels of **(A)** FcγR2a and **(B)** FcγR3a signaling induced by 25 μg/ml of purified IgG from all SARS CoV-2 convalescent donors. Red line represents 2-fold above the mean anti-spike levels of all naïve (CoV-2$^-$) donors in each FcγR signaling assay. Scatter plots show the area under the **(C)** FcγR2a or **(D)** FcγR3a signaling curve versus COVID-19 severity scores. Scatter plots show the area under the **(E)** FcγR2a or **(F)** FcγR3a signaling curve versus the levels of anti-spike IgG titers as quantified by cell-based IgG binding assay. Scatter plots show the area under the **(G)** FcγR2a or **(H)** FcγR3a signaling curve versus the levels of anti-spike IgG titers as quantified by ELISA. All FcγR signaling assays were conducted using a three-point titration curve of purified donor IgG (25ug/ml, 5ug/ml, & 1ug/ml). Area under all points was used to calculate AUC. All FcγR results are representative of at least three different experiments conducted in triplicate. Significance of slopes and $r^2$ values are shown.

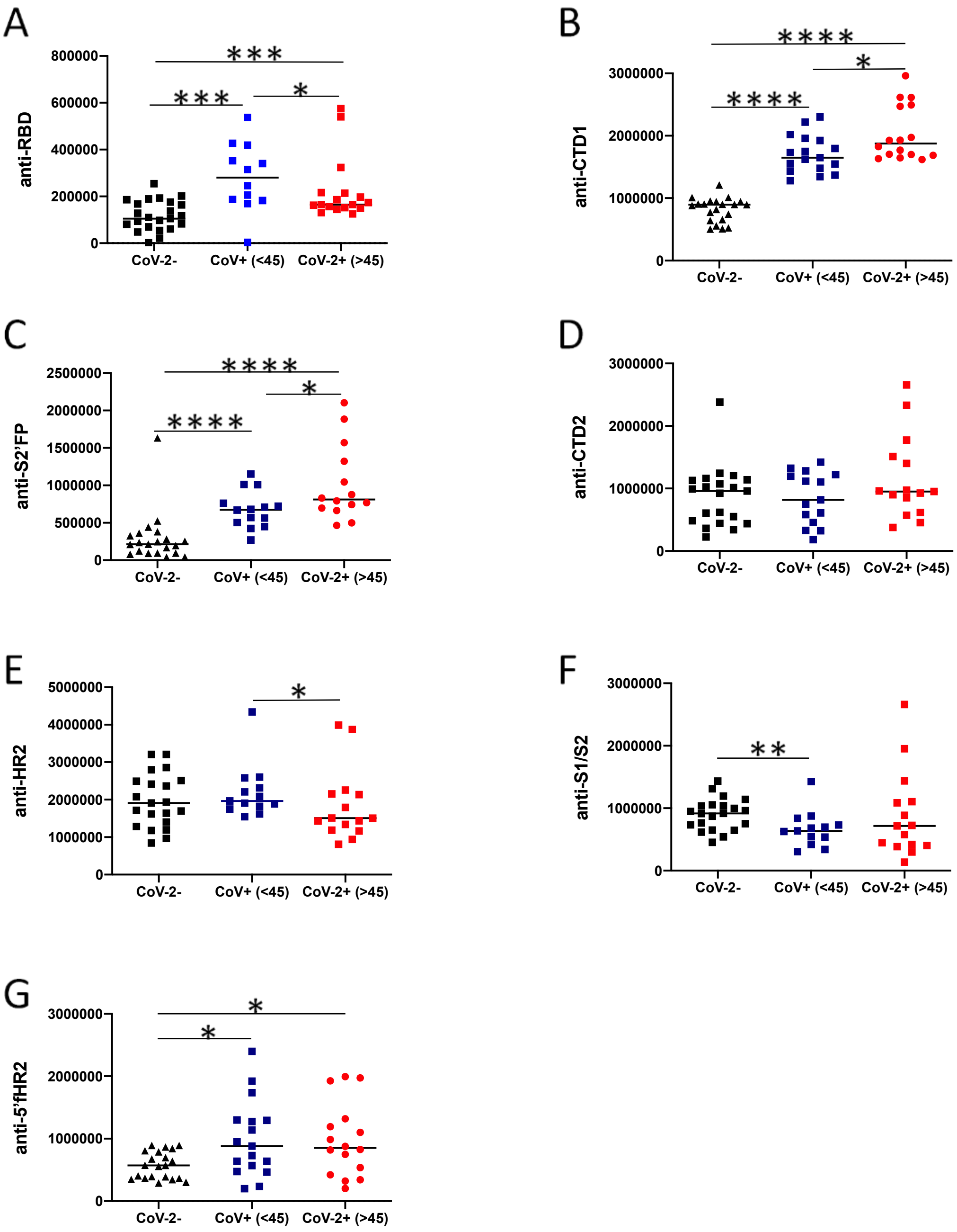


**Supplementary Figure 3. SARS-CoV-2 convalescent IgG differentially target seasonal CoV-conserved and non-conserved SARS-CoV-2 immunodominant epitopes.**

Graphs compare the levels of IgG-binding to **(A)** RBD, (B) CTD1, **(C)** S2'FP, (D) CTD2 **(E)** HR2, (F) S1/S2 **(G)** 5'F HR2 regions in in SARS-CoV-2 naïve (CoV-2-) and SARS-CoV-2 positive convalescent (CoV-2+) donors. Convalescent donors were split into two groups based on COVID19 severity scores, mild (<45), and more severe (>45). The levels of IgG targeting are the mean values obtained from three different experiments conducted in triplicate. Significance was calculated using a 2-tailed Mann-Whitney U test.

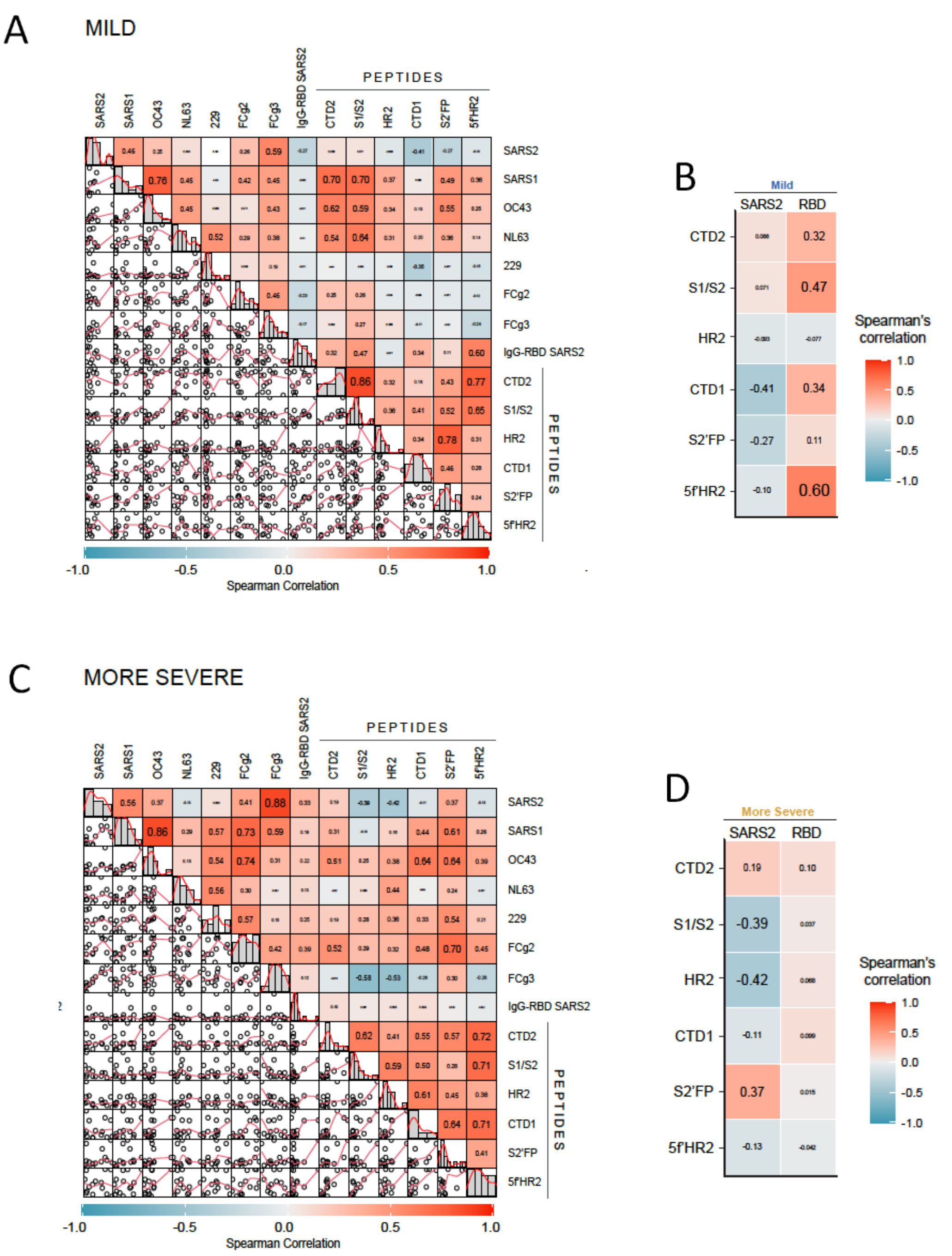


**Supplementary Figure 4. Preferential antibody-binding to different regions of the SARS CoV-2 spike correlates with COVID19 severtiy.**

**(A,C)** Scatter matrix chart summarizes the Spearman's correlation (r values, upper) and the scatter plots (lower) between all analyzed variables for samples separated by mild (n=13) or more-severe (n=15) symptoms, respectively. The small bar graphs (diagonal) represent the distribution of data for each variable. **(B,D)** Heatmaps show the Spearman´s correlations (r values) between IgG-Spike or IgG-RBD and the levels of IgG targeting the six functional spike domains for samples separated by **B)** mild (n=13) or **D)** more-severe (n=15) symptoms.
